# Partially shared representational space for vision and language across objects, scenes, and visual events

**DOI:** 10.64898/2026.11.13.762800

**Authors:** Quan Zou, Kamila M Jozwik

## Abstract

Deep neural networks (DNNs) and behavioural similarity judgements of visual stimuli predict neural responses in visual cortex. Recent work shows that large language model (LLM) embeddings and similarity judgements of language descriptions also predict visual brain activity. These predictors were examined separately across modalities (vision vs language) and types (models vs human behaviour). It remains unclear to what extent these predictors explain shared or unique aspects of visual brain responses, how these contributions vary along the visual hierarchy and category-selective regions, and whether vision-language convergence depends on the stimulus type. We used three fMRI datasets progressing toward naturalistic visual experience, from objects (THINGS) to scenes (NSD) to dynamic events (BMD). For each dataset, we collected similarity judgements of visual stimuli and of their language descriptions from human participants (N = 21-35), extracted vision DNN features and LLM embeddings, and related similarity judgements and model representations to visual brain responses using representational similarity analysis (RSA) and variance partitioning. Vision-language convergence was higher within behaviour than within models. From objects to dynamic events, LLM alignment with vision-based similarity judgements increased, whereas vision DNN alignment remained stable. A shared, predominantly cross-modal component increased along the visual hierarchy and dominated explained variance in high-level visual cortex. LLMs made their largest unique contributions in face– and body-selective regions (FFA and EBA), where shared representations of vision– and language-based human similarity judgements also explained variance beyond the models. In summary, we find a partially shared representational space for vision and language in high-level visual cortex, most clearly in regions representing social information related to people and their actions. As each predictor retained unique variance, combining computational models and human behaviour across vision and language may offer a more comprehensive account of high-level visual representations.

## Introduction

Humans can rapidly make sense of the visual world, whether recognising objects, interpreting scenes, or understanding events. This ability is supported by a hierarchical visual system in which representations become increasingly abstract from early to high-level visual regions (Riesenhuber and Poggio, 1999; DiCarlo et al., 2012). Studies using controlled stimuli and predefined labels have identified high-level regions selective for particular categories, such as faces (Puce et al., 1996; Kanwisher et al., 1997), body parts (Downing et al., 2001; Peelen and Downing, 2005; Bracci et al., 2010) and scenes (Epstein and Kan-wisher, 1998), leading to a view of high-level visual cortex being organised by object category (Grill-Spector and Weiner, 2014).

More recently, studies using naturalistic and large-scale stimulus sets have shown that category-selective regions exhibit multidimensional tuning, capturing finer within-category features and dimensions outside their preferred category, such as food and text (Khosla et al., 2022; van Dyck and Dobs, 2026). The high-level visual cortex has thus been described as a continuous feature space organised along interpretable dimensions (Huth et al., 2012), such as animacy, real-world size, aspect ratio, and semantic content (Huth et al., 2012; Konkle and Caramazza, 2013; Bao et al., 2020; Jozwik et al., 2022; Coggan and Tong, 2023), with growing emphasis on these dimensions’ behavioural relevance (Jozwik et al., 2016; Cichy et al., 2019; Jozwik et al., 2022; Bracci and Op de Beeck, 2023; Contier et al., 2024; Ritchie et al., 2026). For example, beyond their preferred categories, the face-selective fusiform face area (FFA) and the body-selective extrastriate body area (EBA) also carry people– and social-related information, such as sports, social traits, and visual cues of social interaction (Walbrin and Koldewyn, 2019; Abassi and Papeo, 2020; Tsantani et al., 2021; McMahon and Isik, 2023; van Dyck and Dobs, 2026; Yashiro et al., 2026).

These dimensions cover both visual and conceptual properties (Bracci and Op de Beeck, 2016; Jozwik et al., 2016; Hebart et al., 2020; Jozwik et al., 2023; Contier et al., 2024). Representations become increasingly abstract and conceptual along the ventral and lateral visual pathways towards the anterior border of visual cortex (Lingnau and Downing, 2015; Devereux et al., 2018; Wurm and Caramazza, 2022; McMahon et al., 2023). Several lines of evidence suggest that these conceptual representations are encoded similarly for visual and language input. Semantic maps estimated from movies and from stories align (Popham et al., 2021), encoding models trained on responses to stories predict responses to movies and vice versa (Tang et al., 2023), and a single decoder can identify stimuli presented as either images or captions (Nikolaus et al., 2026). These findings motivate the hypothesis that vision and language converge on a shared high-level representational space (Doerig et al., 2025; Simkova et al., 2026).

Characterising this representational space also requires anchoring it in behaviour, since cortical representations are shaped by the behavioural goals that vision serves (Bracci and Op de Beeck, 2023; Ritchie et al., 2026). Similarity judgements reflect the multidimensional mental representations of objects (Hebart et al., 2020), revealing dimensions that predict responses across visual cortex better than conventional object categories (Contier et al., 2024). Judgements of visual stimuli and of their language descriptions also overlap substantially (Dima et al., 2024; Simkova et al., 2026) and explain largely shared variance in high-level visual cortex (Simkova et al., 2026), suggesting that concepts are organised similarly across vision and language.

Whereas similarity judgements reveal how people mentally organise concepts, computational models trained on a single modality can test whether this shared structure can be learned from vision or language alone (Conwell et al., 2025). Deep neural networks (DNNs) trained on computer vision tasks have served as models of visual cortex, capturing its hierarchical progression from low-to high-level features (Khaligh-Razavi and Kriegeskorte, 2014; Yamins et al., 2014; Yamins and DiCarlo, 2016). Large language models (LLMs) trained on large text corpora have similarly served as models of language processing in the brain (Schrimpf et al., 2021; Goldstein et al., 2022). Recent work shows that LLM embeddings also predict responses in high-level visual cortex (Lahner et al., 2024; Doerig et al., 2025), likely because training on text allows models to acquire conceptual relations and world knowledge that also structure visual representations (Petroni et al., 2019; Roberts et al., 2020). These findings are consistent with the Platonic Representation Hypothesis, which proposes that models trained on different modalities increasingly converge on a shared statistical model of the world (Huh et al., 2024).

Vision-language convergence has thus been reported in the brain, in behaviour, and in models, but largely in separate studies that differ in stimuli, predictors, and analyses (e.g., Dima et al., 2024; Garcia et al., 2025; Simkova et al., 2026), making it difficult to tell whether these findings reflect the same underlying structure. Three questions therefore remain unanswered. First, it is unclear to what extent the behavioural and computational predictors of vision and language explain shared or distinct aspects of visually evoked brain responses. Second, little is known about how these contributions vary along the visual hierarchy and across category-selective regions, which differ in the dimensions they represent and in their level of abstraction. Third, it has not been tested whether vision-language convergence and the contribution of each predictor depend on the type of visual stimulus. Compared with object images, scenes contain multiple objects and their spatial and contextual relations (Henderson and Hollingworth, 1999; Epstein, 2005; Võ, 2021), and dynamic events further add agents and actions unfolding over time (Zacks, 2020; Lahner et al., 2024). These stimulus types thus carry different kinds and amounts of information, which language may capture to different degrees. Comparing the stimulus types within a single framework can reveal how the contributions of vision and language, and of behaviour and models, change as visual input becomes more complex and naturalistic.

In this study, we use three fMRI datasets that form a progression towards natural visual experience, from static objects (THINGS-fMRI; Hebart et al., 2023) to static scenes (Natural Scenes Dataset; Allen et al., 2022) and dynamic events (BOLD Moments Dataset; Lahner et al., 2024). For each dataset, we collected similarity judgements of the visual stimuli and of their language descriptions, and extracted representations from vision DNNs and LLMs. Using representational similarity analysis (RSA), we first examined how these predictors relate to one another and to brain responses, and then used variance partitioning to separate their shared and unique contributions in retinotopically defined early to mid-level visual areas and functionally defined category-selective regions. These analyses allowed us to test to what extent vision– and language-derived predictors explain shared or distinct variance in visually evoked brain responses. In particular, we examined whether language contributes unique explained variance beyond vision, and whether human behavioural similarity judgements contribute unique explained variance beyond computational models. We further asked how these contributions vary along the visual hierarchy and across category-selective regions, and whether they differ across objects, scenes, and dynamic events.

## Methods

### Functional MRI Datasets

The present study uses three large-scale fMRI datasets, each containing responses to a different class of visual stimuli: objects (THINGS-fMRI; Hebart et al., 2023), natural scenes (Natural Scenes Dataset; Allen et al., 2022), and visual events (BOLD Moments Dataset; Lahner et al., 2024). From each dataset, we select a subset of approximately 100 stimuli (100 for THINGS and BMD, 102 for NSD; see Figure 1a), together with their corresponding fMRI responses. These subsets were selected because they were either shared across participants and sessions, had higher repetition counts, or both, making them well suited for representational similarity analysis. For THINGS-fMRI and BMD, these subsets correspond to the respective test sets used for model evaluation. For NSD, we use the designated Special100 subset, curated by the dataset authors to provide broad coverage of the semantic space for a Multiple Arrangement Task. Across all three datasets, we used single-trial beta estimates obtained using GLMsingle (NSD, BMD) or ICA-denoised GLM (THINGS; see Hebart et al., 2023 for details), and predefined regions of interest (ROIs) covering early visual and category-selective areas together with their respective noise-ceiling (NC) estimates, all provided by each dataset. Brief summaries of each dataset and the components relevant to the present study are provided below, followed by a description of the standardisation procedure used to facilitate cross-dataset comparison. For further details, please see the original dataset publications (Allen et al., 2022; Hebart et al., 2023; Lahner et al., 2024).

**Fig. 1.**
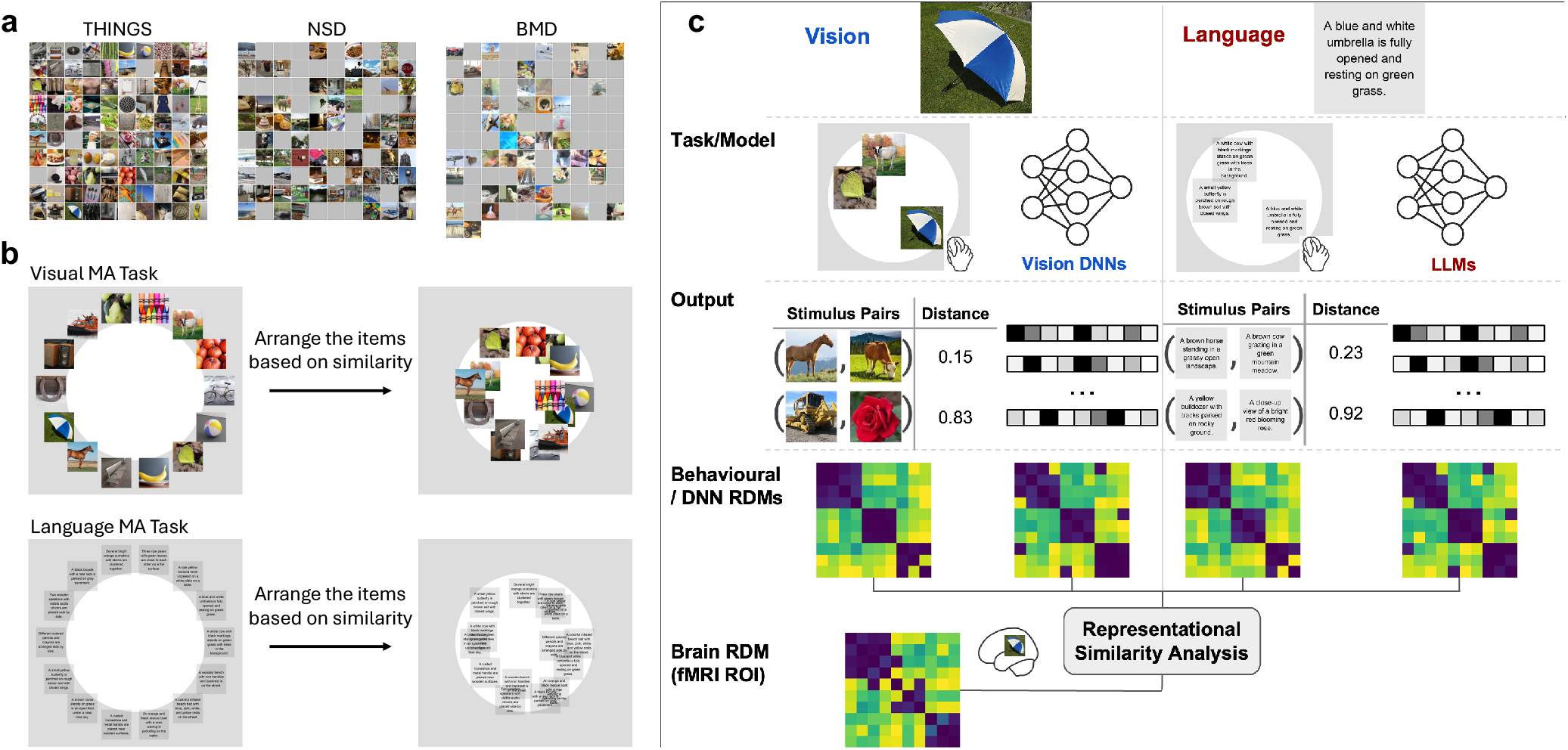
Stimulus sets, similarity judgement tasks, and analysis pipeline. ***a***, Stimulus sets. The 100 object images (THINGS), 100 scene images (NSD), and 102 video clips (BMD; one representative frame shown per clip) were used in the analyses. Grey squares are placed over face-containing images in line with bioRxiv policy. ***b***, Multiple Arrangement (MA) tasks. On each trial, a subset of stimuli appeared around a circular arena. Participants were instructed to drag and place similar stimuli closer together and dissimilar stimuli further apart, so that on-screen distance reflected perceived dissimilarity (left, trial onset; right, completed arrangement). Top, visual MA task using images or video clips; bottom, language MA task using the corresponding sentence descriptions. The two task variants were procedurally identical apart from the stimulus type, so any difference between the resulting RDMs reflects the modality of the judgements rather than task demands. Across trials, stimulus subsets were adaptively sampled to estimate the full pairwise dissimilarity structure efficiently, and stimuli that were inconsistently judged between trials were sampled more on subsequent trials to obtain high-consistency judgements. On-screen distances were combined by inverse multidimensional scaling to yield one representational dissimilarity matrix (RDM) per participant. ***c***, Analysis pipeline. For each visual stimulus and its language description, two parallel processing streams were applied to derive a vision-based representation (left, blue) and a language-based representation (right, red). Task/Model: Behavioural similarity judgements were collected from human participants using the MA tasks in two variants: arranging the images or videos (vision) or arranging their short sentence descriptions (language). In parallel, representations were extracted from pretrained vision DNNs (image-input DNNs for images, video-input DNNs for videos) and LLMs. Output: The MA tasks produced pairwise on-screen distances between stimuli, reflecting perceived dissimilarity. The vision DNNs and LLMs produced layer-wise activation vectors per stimulus. Behavioural/DNN RDMs: Pairwise distances (behaviour) and pairwise distances between activation vectors (models) were assembled into RDMs, producing four RDMs per stimulus set: vision behaviour, vision DNN, language behaviour, and LLM. Brain RDM: A brain RDM was computed per fMRI region of interest (ROI) from the pairwise distances between stimulus-evoked response patterns. Each behavioural and model RDM was then compared with each brain RDM via RSA, enabling direct cross-system and cross-modal comparison of representational geometry. The pipeline was applied identically to all three fMRI datasets.

#### THINGS-fMRI

The THINGS dataset contains fMRI responses from three participants who viewed 8,740 unique images spanning 720 object concepts, acquired using whole-brain 3T fMRI at 2 mm isotropic resolution (TR = 1.5 s). The object concepts include living and non-living objects, non-countable substances, faces, and body parts, with each concept represented by 12 images of the object embedded in a naturalistic background (Hebart et al., 2019). During scanning, the participants reported the presence of catch images (images depicting unrecognisable novel objects created using generative adversarial neural network BigGAN, see Brock et al., 2019) using a button press. The present analysis focuses on the 100 test images, each of which was presented once per session across all 12 sessions (100 *×* 12 = 1200 trials).

#### Natural Scenes Dataset (NSD)

The NSD consists of fMRI responses from 8 participants to 73,000 unique natural scene images, acquired with whole-brain 7T fMRI at 1.8 mm isotropic resolution (TR = 1.6 s). The images were drawn from the Microsoft Common Objects in Context (COCO) dataset (Lin et al., 2014), with each image accompanied by five human-written captions (Chen et al., 2015). During scanning, the participants performed a long-term continuous recognition task, judging whether each image was novel or previously presented using a button press. For the present study, we use the Special100 subset (Allen et al., 2022), in which each stimulus was presented three times per participant (100 *×* 3 = 300 trials), and the 1.8-mm volume preparation of NSD data (betas_fithrf_GLMdenoise_RR).

#### BOLD Moments Dataset (BMD)

The BMD consists of fMRI responses from 10 participants to 1102 naturalistic 3-s video clips of visual events, acquired using whole-brain 3T fMRI at 2.5 mm isotropic resolution (TR = 1.75 s, resampled to 1 s). The videos were drawn from the Memento10k dataset (Newman et al., 2020) and each video was accompanied by five human-generated text descriptions obtained via Amazon Mechanical Turk (AMT) and provided as part of BMD (Lahner et al., 2024). During scanning, the participants reported the presence of null trials (grey screen) using a button press. The present analysis focuses on the 102 test videos, each presented 10 times per participant (102 *×* 10 = 1020 trials). We used the version B preprocessed data provided by the dataset authors (Lahner et al., 2024). These data contain GLMsingle-derived single-trial beta estimates, making them comparable to the corresponding THINGS and NSD preprocessed data (Allen et al., 2022; Hebart et al., 2019).

#### Cross-Dataset Standardisation

The three datasets differ in their predefined ROI schemes and in the form of their available noise-ceiling estimates. To enable reliable comparisons, we applied the following steps to better align the three datasets. First, we restricted cross-dataset analyses to functionally defined ROIs available in all three datasets. In each dataset, ROIs were defined for each participant using independent retinotopic mapping or pRF scans, along with category-selective functional localizers contrasting faces, body parts, scenes, words, and objects. These procedures provided early-to mid-visual and functionally defined category-selective ROIs in each participant’s native space. We retained ten ROIs common to all three datasets: early to mid-visual areas V1, V2, V3, and hV4, extrastriate body area (EBA), occipital face area (OFA), fusiform face area (FFA), occipital place area (OPA), parahippocampal place area (PPA), and retrosplenial cortex (RSC), see Figure 2. Second, we selected voxels with reliable stimulus-driven response patterns by applying a common noise ceiling (NC) threshold across datasets, where NC is the percentage of variance in the trial-averaged responses that is explainable by the stimulus. The NC estimates follow the analytical framework described by Allen et al. (2022), in which per-voxel beta variance is decomposed into signal and noise components. This analysis yields a noise-ceiling signal-to-noise ratio, defined as NCSNR = *σ*_signal_*/σ*_noise_, which can be converted to the percentage of explainable variance as:

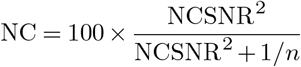

where *n* is the number of repetitions averaged per condition. NSD provides NCSNR values, which we converted using *n* = 3 for the shared test-set images. BMD Version B and THINGS-fMRI provide NC estimates directly for the test sets. We retained voxels with NC *>* 10% to exclude voxels with weak stimulus-driven responses and to reduce differences in data reliability. This threshold approximates the NCSNR *>* 0.2 cutoff used by Conwell et al. (2024) for NSD, which corresponds to NC *≈* 10.7% when *n* = 3. Third, following the normalisation procedure recommended by Allen et al. (2022) and adopted in subsequent work (Doerig et al., 2025; Gifford et al., 2023), we applied voxel-wise z-scoring per session to the single-trial beta estimates in THINGS and NSD. We subtracted each voxel’s session-wise mean and divided it by its session-wise standard deviation. BMD provides single-trial betas that are already normalised, so no further normalisation was applied. Finally, for each participant and ROI, we extracted z-scored single-trial beta estimates from all voxels within the ROI and averaged beta estimates across repetitions of each stimulus, which resulted in one response pattern per stimulus per ROI.

**Fig. 2.**
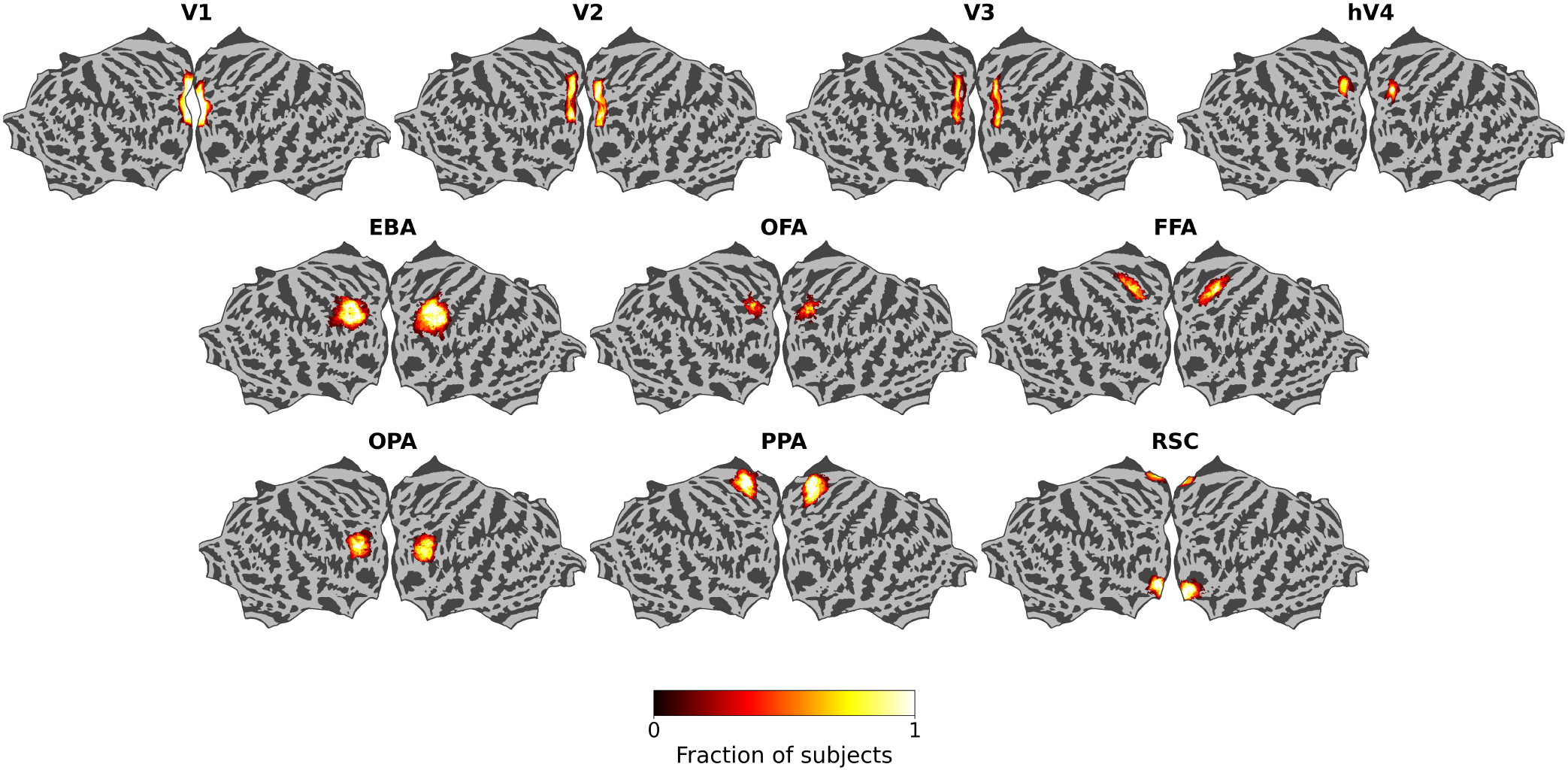
Anatomical locations of the regions of interest. Probabilistic maps of the early to mid-visual areas (V1, V2, V3, hV4) and the functionally defined category-selective regions EBA, OFA, FFA, OPA, PPA, and RSC, shown on the flattened fsaverage surface. Maps are taken from the NSD dataset (Allen et al., 2022), in which each manually defined ROI was mapped to fsaverage by nearest-neighbour interpolation and the fraction (between 0 and 1) of participants with that ROI present was computed at each vertex; colour indicates this fraction. Here, V1, V2, and V3 combine the NSD-released ventral and dorsal subdivisions, and FFA combines FFA-1 and FFA-2. This figure is for illustration only, to show the approximate cortical locations of the ROIs. All analyses were carried out in volume space: in each participant’s native space for THINGS and NSD, and in MNI152NLin2009cAsym for BMD.

### Behavioural Similarity Judgement

To characterise the representational structure underlying vision and language behaviour, we collected behavioural similarity judgements (Mur et al., 2013) in each modality.

#### Participants

Forty-three healthy adults (11 females, mean age = 29.9 years, SD = 7.4 years) were recruited from Prolific (`www. prolific.com`) and completed behavioural similarity judgements across one or more of three stimulus sets. Participant numbers per stimulus set were as follows: THINGS (visual, N = 25; language, N = 28; both, N = 23), NSD (visual, N = 23; language, N = 25; both, N = 15), and BMD (visual, N = 21; language, N = 35; both, N = 20). Thirty-seven participants completed similarity judgements for more than one dataset (two datasets, N = 18; all three datasets, N = 19). Six participants completed similarity judgements for one dataset only. All participants provided informed consent and were compensated for their time. The experiments were approved by the Institute for Technology and Humanity Ethics Committee at the University of Cambridge.

#### Stimuli

For the visual condition, we presented the original images or video clips from each stimulus set. For the language condition, we used human-generated captions describing the corresponding visual stimuli. NSD and BMD each provide five human-written captions per stimulus; for THINGS, we collected an analogous set of five captions per stimulus from a separate group of participants in a labelling task. For each stimulus, GPT-4.1 mini was used to synthesise a single representative caption from the five human annotations (for full prompting instructions, see “Stimuli and caption generation” in the Supplementary Information). To approximate the visual format of the image stimuli, each caption was rendered as centred text on a 500 *×* 500 pixel canvas with a semi-transparent grey background.

#### Multiple Arrangement Task

We collected pairwise similarity judgements using the multiple arrangement (MA) task (Mur et al., 2013) administered online via the Meadows platform (www.meadows-research.com). For each dataset, participants first completed the language MA task, followed by the visual MA task. The language task was always administered first to minimise visual bias in language-based judgements. Participants spent 113 minutes per task on average, with the language task (mean = 120 min) taking significantly longer than the visual task (mean = 105 min; Wilcoxon signed-rank test, *p <* 0.001). On each trial, stimuli (either images/video clips or rendered captions) were displayed around a white circular arena, and participants were instructed to arrange stimuli inside the arena based on their similarity, placing similar items close together and dissimilar items further apart (Figure 1b). To avoid biasing participants towards a particular feature dimension, instructions did not specify which properties of the stimuli to attend to.

### Model Representations

To characterise vision and language representational structures of the models, we extracted models’ hidden-layer activations from vision Deep Neural Networks (DNNs) and Large Language Models (LLMs).

#### Model Selection

We selected architecturally diverse models from two modalities (vision and language), with vision DNNs additionally matched to each dataset’s stimulus format (image-input or video-input). For each dataset, we used four vision and four language models to enable balanced cross-modal comparisons.

### Vision DNNs

For static images in THINGS and NSD, we used four image-input vision models: AlexNet (Krizhevsky et al., 2012), VGG16 (Simonyan and Zisserman, 2015), ResNet50 (He et al., 2016), and ViT-B-P16 (Dosovitskiy et al., 2021). AlexNet served as a canonical DNN baseline model commonly used in model-brain alignment studies (Khaligh-Razavi and Kriegeskorte, 2014; Cichy et al., 2016; Jozwik et al., 2017). VGG16 represented a deeper DNN architecture, whereas ResNet50 introduced residual connections while keeping a standard visual backbone. The ViT-B-P16 (vit_base_patch16_224) represents the transformer-based vision paradigm, replacing convolutional inductive biases with patch-based self-attention. AlexNet, VGG16, and ResNet50 were pretrained on ImageNet-1k for supervised image classification. ViT-B-P16 was pretrained on ImageNet-21k under the same classification objective and fine-tuned on ImageNet-1k, allowing representations learned at scale to be adapted to the specific ImageNet-1k benchmark and making the vision DNNs more comparable (Dosovitskiy et al., 2021). Together, these models span major architectural paradigms in static visual object recognition, from canonical DNNs through residual ones to vision transformers.

For dynamic visual events in BMD, we used four video-input vision models: TSM-R50 (Lin et al., 2019), SlowFast-R50 (Feichtenhofer et al., 2019), VideoMAE-B (Tong et al., 2022), and TimeSformer-divST (Bertasius et al., 2021). TSM-R50 augments a 2D ResNet backbone with temporal shift operations for lightweight temporal mixing. SlowFast-R50 uses dual 3D-DNN pathways that operate at different frame rates to capture slow semantic and fast motion information. VideoMAE-B is a self-supervised video transformer trained with masked reconstruction over space-time tube embeddings. TimeSformer-divST is a supervised video transformer that factorises self-attention into temporal and spatial operations. All four models are pretrained on Kinetics-400 via supervised classification, except VideoMAE-B, which is first self-supervised by masked-video reconstruction and then fine-tuned on standard video action recognition benchmarks (Tong et al., 2022). Together, these models span supervised learning and self-supervised pretraining across DNN and transformer backbones, with distinct temporal modelling strategies.

### LLMs

For all three datasets, we used four language models: GPT-2 (Radford et al., 2019), BERT (bert-base-uncased; Devlin et al., 2019), MPNet (all-mpnet-base-v2; Song et al., 2020), and MiniLM-L6 (all-MiniLM-L6-v2; Wang et al., 2020). GPT-2 and BERT were included as token-level contextual language models: GPT-2 is an autoregressive transformer trained with left-to-right language modelling, whereas BERT is a bidirectional transformer trained with masked language modelling. MPNet and MiniLM-L6 were included as sentence-embedding models optimised for semantic similarity: MPNet combines masked and permuted language modelling objectives, whereas MiniLM-L6 is a distilled sentence-embedding model that provides a compact semantic representation. GPT-2 and BERT were pretrained as general-purpose language models; MPNet and MiniLM were additionally fine-tuned on over 1 billion sentence pairs with a contrastive sentence-similarity objective (Song et al., 2020; Wang et al., 2020).

#### Feature Extraction

We accessed all models (except the sentence LLMs) and extracted model features using the Net2Brain toolbox (Bersch et al., 2025), a Python package for comparing DNN representations with brain responses. For vision models, Net2Brain applied each model’s source-library default preprocessing to each stimulus and extracted features from a curated set of layers. We extracted the output of the successive processing stages across the full depth of each DNN. For image-input models, features were extracted from eight AlexNet layers comprising five convolutional and three fully connected layers; eight VGG16 layers comprising five block-end max-pooling outputs and three fully connected layers (Ye et al., 2025); five ResNet50 layers comprising the four backbone stage outputs and the final global average-pooling layer; and all 12 transformer blocks of ViT-B-P16. For video-input models, with per-model frame sampling, features were extracted from the four backbone stages of TSM-R50, matching the BMD encoding analysis (Lahner et al., 2024); the slow and fast pathways of SlowFast-R50; and all 12 transformer blocks of VideoMAE-B and TimeSformer-divST. For contextual token-level language models (GPT-2 and BERT), each caption was tokenised and passed through the model to obtain token-wise activations at each layer. We used last-token pooling for GPT-2 (Schrimpf et al., 2021) and mean pooling for BERT (Toneva and Wehbe, 2019), and included all 12 transformer layers (each 768-dimensional). For sentence-embedding language models (MPNet and MiniLM), features were extracted outside Net2Brain using a dedicated sentence-level pipeline. Captions were tokenised with padding and truncation to a maximum length of 128 tokens, passed through the model, and mean-pooled over the final hidden state using the attention mask to obtain one embedding per caption (768-dimensional for MPNet, 384-dimensional for MiniLM), following previous work (Doerig et al., 2025).

### Representational Similarity Analysis (RSA)

We performed representational similarity analysis (RSA; Kriegeskorte et al., 2008a) using the Python package *rsatoolbox* (van den Bosch et al., 2025), which provides tools for representational dissimilarity matrix (RDM) construction, model comparison, and noise-ceiling estimation.

#### Construction of Representational Dissimilarity Matrices

For each dataset, we constructed RDMs based on brain responses, behaviour, and model representations (Figure 1c). Brain RDMs were computed per participant and ROI as pairwise distances between voxel response patterns. For each participant, a behavioural RDM was estimated from the on-screen distances between stimuli in the MA arena after the distances were refined across trials (Kriegeskorte and Mur, 2012). Per-participant RDMs were averaged to obtain group-level vision and language RDMs per dataset, following Simkova et al. (2026). DNN RDMs were constructed from extracted layer activations using Net2Brain’s RDMCreator: for each selected layer of each model, pairwise distances between stimulus activation vectors produce an RDM. Following previous work, we used Pearson’s correlation distance (1 *− r*) to create brain and model RDMs (Conwell et al., 2024; Doerig et al., 2025), and Euclidean distance to create behavioural RDMs, as this data was derived from spatial arrangement via inverse multidimensional scaling (MDS) (Simkova et al., 2026; van den Bosch et al., 2025). All RDMs were stored as the vectorised upper triangle over all stimulus pairs.

#### Alignment Analysis

We assessed representational similarity between behavioural RDMs, model RDMs, and the brain RDMs (Figure 1c) using Pearson’s correlation. We used *rsatoolbox*’s `eval_fixed` function (van den Bosch et al., 2025). Each comparison produced one correlation per participant, which we summarised across participants as the group mean. To assess model-behaviour alignment, we compared the model RDMs with each participant’s behavioural RDM. Model RDMs were defined as the best layer within each model family, selected separately for each dataset and similarity judgement task by leave-one-participant-out cross-validation. For each held-out participant, we selected the layer with the highest mean correlation across the remaining participants, and scored alignment as its correlation with the held-out participant’s RDM. To assess behaviour-brain and model-brain alignment, we compared behavioural and model RDMs with each fMRI participant’s brain RDM for each ROI. Behavioural RDMs were the group-mean vision-based and language-based similarity judgement RDMs of each dataset. Model RDMs were defined as the best layer within each model family, selected separately for each ROI as the layer most often selected across 10 random 70:30 stimulus splits. In each split, we selected the layer with the highest mean correlation across fMRI participants on the training stimuli, and we used the layer selected most often across splits. Alignment of this layer was then computed using all stimuli.

#### Noise Ceiling Estimation

Noise ceilings (NCs) were estimated from participants’ vision– and language-based behavioural RDMs for each dataset, and from participants’ brain RDMs for each ROI in each dataset using Pearson’s correlation (Nili et al., 2014), as implemented in the *rsatoolbox* (van den Bosch et al., 2025). The lower bound is the mean leave-one-out correlation between each participant’s RDM and the average RDM of the other participants. The upper bound is the mean correlation between each participant’s RDM and the group-mean RDM (including that participant).

### Variance Partitioning

We quantified shared and unique contributions to brain alignment via variance partitioning with cross-validated non-negative least-squares (NNLS) regression (Jozwik et al., 2016; Simkova et al., 2026). For each participant, ROI, and predictor pair (A, B), we fit three NNLS models (A alone, B alone, and joint A + B), each scored by cross-validated *R*^2^ over 10 random 70:30 stimulus splits. Averaging across folds, we computed Unique A contribution 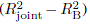, Unique B contribution 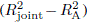, and shared contribution 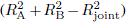. We performed four pairwise partitions and one four-way partition. The pairwise partitions compared are: (1) vision-versus language-based behavioural similarity judgement RDMs; (2) best vision-DNN layer versus best language-model layer RDMs, selected per ROI by nested cross-validation; (3) vision DNNs versus vision-based similarity judgement RDMs; and (4) LLMs versus language-based similarity judgement RDMs. The first two partitions assessed the relative contributions of vision and language within behaviour and models, respectively, whereas the latter two assessed the unique and shared contributions of behaviour and models within each modality. The four-way partition extends this analysis to all four predictor RDMs, crossing modality (vision, language) with system (human behaviour, model): vision behaviour (*B_V_*), vision DNN (*M_V_*), language behaviour (*B_L_*), and LLM (*M_L_*). For each participant and ROI, we fitted all 15 non-empty subsets of the four predictors (4 single predictors, 6 pairs, 4 triples, and the full model) by NNLS on the same 10 stimulus-level 70:30 splits. Each predictor’s unique contribution is the drop in *R*^2^ when that predictor is removed from the full model 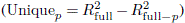. The shared contribution is the remaining variance explained jointly by two or more predictors 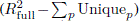. The four-way partition thus asks what each predictor adds beyond the other three, rather than beyond a single predictor. To quantify how much of the shared variance is cross-modal, we treated the vision predictors (*B_L_*, *M_L_*) and the language predictors (*B_L_*, *M_L_*) as two blocks and computed their shared variance as in the pairwise partitions 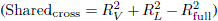. Here, 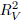 and 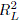 are the *R*^2^ of the vision-only and language-only models. This cross-modal component includes all shared variance except the variance shared only within one modality. Because all components are differences between cross-validated *R*^2^ values, they can be negative even though NNLS constrains regression weights to be non-negative. Negative values were retained for statistical testing but set to zero for visualisation and reported percentages. We additionally performed two pairwise partitions, comparing (1) behavioural versus DNN predictors and (2) vision-versus language-derived predictors, by combining predictors within each source or modality, respectively. These analyses are reported in the Supplementary Information.

## Statistical Analysis

### Model-Behaviour Alignment Statistical Analysis

For each model, we correlated its best-fitting layer with every participant’s behavioural RDM using Pearson’s correlation, giving one alignment value per participant. We then tested across participants whether each model aligned with behaviour above zero (two-sided Wilcoxon signed-rank test), and whether the best vision DNN layer and the best LLM layer differed in how well they aligned with behaviour (paired two-sided Wilcoxon signed-rank test, same participants). To test for trends across stimulus types, we applied two-sided Jonckheere-Terpstra tests (Jonckheere, 1954) to the alignment values ordered from objects (THINGS) to scenes (NSD) to dynamic events (BMD). All *p*-values were FDR-corrected (Benjamini-Hochberg, *α* = 0.05) within each dataset.

### Behaviour and Model Brain Alignment Statistical Analysis

For each ROI, RSA produced one alignment score with the brain RDM for both behavioural and model RDMs per participant. The scores were computed using Pearson’s correlation, and two group-level tests were applied. First, to assess whether each predictor RDM aligned with the brain RDM above chance, the alignment scores were tested against zero. Second, to compare predictor alignment within each ROI, per-participant differences were tested against zero. We used two-sided Wilcoxon signed-rank tests. For THINGS (N = 3), the Wilcoxon test was underpowered, so both tests were instead implemented with two-sided stimulus-label permutation tests on the group-mean statistics (Kriegeskorte et al., 2008b). For each ROI, the stimulus labels of all participants’ brain RDMs were jointly shuffled 1,000 times to build null distributions. The inference therefore generalises across stimuli rather than participants. A direction (e.g., vision *>* language or vice versa) was reported only when the difference was significant and the better-aligned predictor exceeded zero. All *p*-values were FDR-corrected across the ROIs (Benjamini-Hochberg, *α* = 0.05) within each dataset.

### Variance Partitioning Statistical Analysis

All partition statistics were computed on cross-validated *R*^2^. For each ROI, unique and shared components were tested against zero. In the four-way partition, predictors were additionally compared pairwise on the variance each explained alone. As in the alignment analysis, we used two-sided Wilcoxon signed-rank tests across participants, replaced for THINGS by stimulus-label permutation tests. All *p*-values were FDR-corrected (Benjamini-Hochberg, *α* = 0.05) across ROIs within each dataset.

## Results

To interpret how vision– and language-derived predictors capture the representational space of visual cortex, we first characterised the predictors before relating them to brain data. We examined the reliability of similarity judgements and the overlap between predictor modality (language-vs vision-based) and predictor type (model representations vs human behaviour), across three datasets progressing towards more naturalistic visual experience. We then measured how well the predictors explained visual cortex representations and examined the shared and unique explained variance, exploring both the hierarchy of low-to high-level visual areas and the heterogeneity of category-selective regions.

### Vision-language convergence is strong within behaviour and weaker within models

Before testing whether vision– and language-based similarity judgements converge, we assessed how consistently participants agreed with one another within each modality’s similarity judgements. We estimated the within-modality reliability for the vision– and language-based multiple-arrangement (MA) tasks using leave-one-out inter-participant Pearson’s correlations (Nili et al., 2014). Reliability was significantly above zero in every dataset and modality (two-sided Wilcoxon signed-rank tests, all FDR-corrected *p <* 0.001; Figure 3a), and increased from THINGS (objects) to NSD (scenes) to BMD (video clips) for both vision (*r* = 0.29, 0.32, 0.45) and language behaviour (*r* = 0.13, 0.26, 0.35). Language reliability was numerically lower than vision reliability in all datasets, and significantly so in THINGS and BMD (two-sided Mann-Whitney test, FDR-corrected *p <* 0.05), indicating that language-based similarity judgements were less consistent across participants than vision-based similarity judgements. The language MA task also took longer to complete than the vision MA task (mean = 120 vs 105 min, respectively; two-sided Wilcoxon signed-rank test, *p <* 0.001), possibly because participants had to infer what each description depicted, which can differ across individuals and lead to lower reliability.

**Fig. 3.**
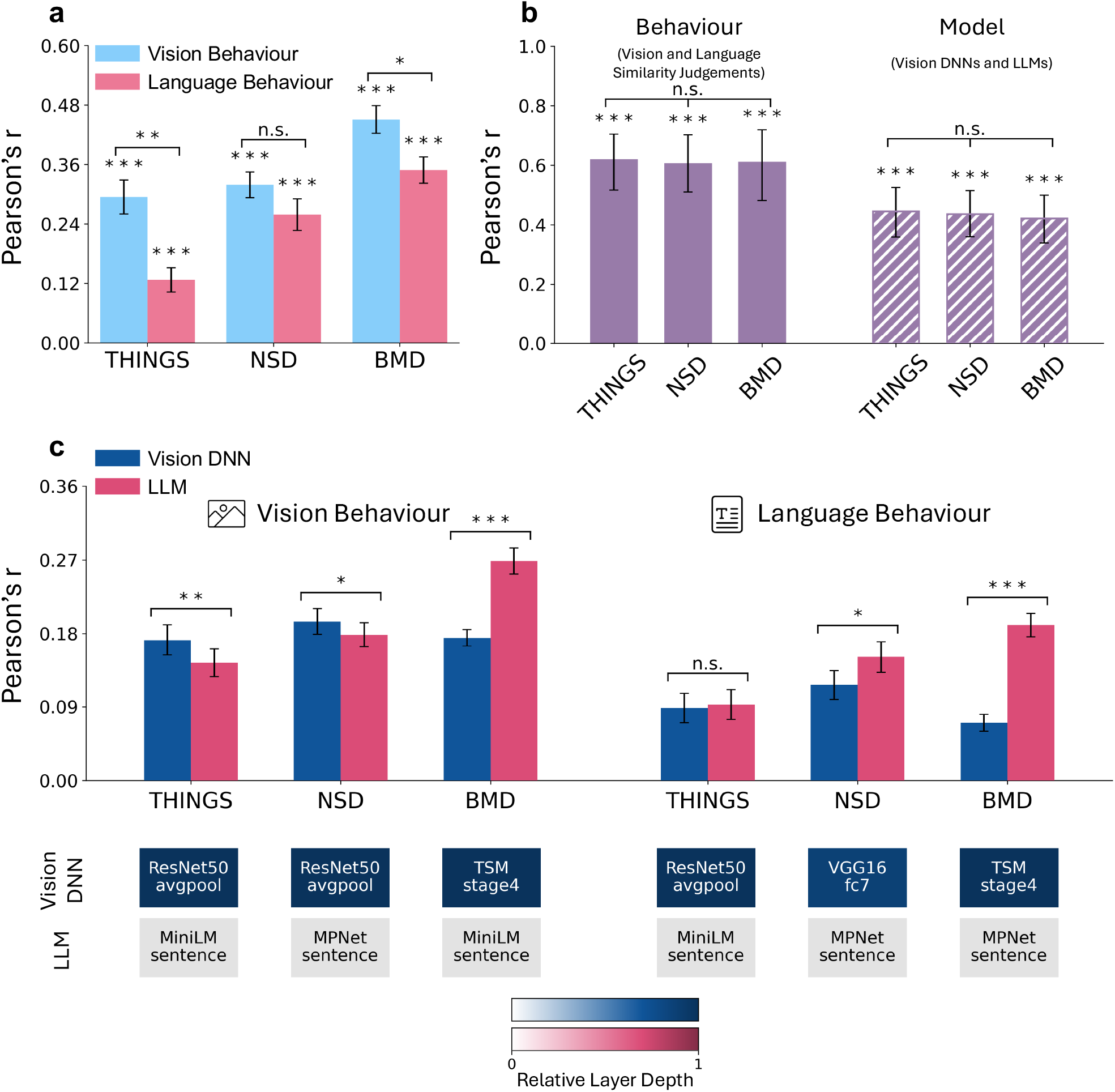
Vision-language correspondence in behaviour and models. ***a***, Within-modality inter-participant reliability of vision and language behaviour. Inter-participant reliability is quantified as Pearson’s *r*. Error bars indicate SEM across participants. Asterisks above bars denote reliability significantly greater than zero (two-sided Wilcoxon signed-rank tests). Brackets indicate vision-language differences (two-sided Mann-Whitney U tests). ***b***, Correlation between vision and language RDMs for behaviour (vision– and language-based similarity judgements) on the left, and for models (vision DNNs and LLMs) on the right, in each dataset (THINGS, NSD, BMD). Behavioural RDMs are averaged at the group level. For each model, the layer that best matches the same-modality judgements (vision-based for vision DNNs, language-based for LLMs) is selected, and the RDMs of the selected layers are averaged within each model family to obtain one vision and one language model RDM per dataset. Vision and language RDMs for behaviour and model are then correlated using Pearson’s correlation. Asterisks above bars denote correlations significantly greater than zero (two-sided stimulus-label permutation tests). Brackets indicate the significance of between-dataset comparisons based on stimulus bootstrapping. ***c***, Model-behaviour alignment. Bars show how well the vision DNNs and LLMs capture human vision-based (left three groups) and language-based (right three groups) similarity judgements in each dataset. For each model family, dataset, and judgement task, the model layer was selected by leave-one-participant-out cross-validation. Alignment is the Pearson’s correlation between the selected layer’s RDM and the held-out participant’s behavioural RDM, averaged across participants. Alignment relative to the noise ceiling is provided in Supplementary Figure 2. All alignments were significantly above zero. Brackets denote the vision DNN versus LLM difference in explaining the similarity judgements. We used two-sided Wilcoxon signed-rank tests. All *p*-values were FDR-corrected. Significance: \**p <* 0.05, \*\**p <* 0.01, \*\*\**p <* 0.001. Below each pair of bars, the selected vision DNN (upper row) and LLM (lower row) are shown, shaded by relative layer depth (layer position divided by the number of extracted layers, 1 = final layer). MPNet and MiniLM provide a single sentence embedding and are shown in grey.

Given that similarity judgements in both modalities were reliable, we next quantified the convergence between vision– and language-based similarity judgements within each dataset (Figure 3b, left). The group-average vision– and language-based similarity judgements RDMs were highly correlated in all three datasets (THINGS: Pearson’s *r* = 0.62; NSD: *r* = 0.60; BMD: *r* = 0.61; two-sided stimulus-label permutation tests, all FDR-corrected *p <* 0.001), with no significant differences across datasets (stimulus bootstrap; all pairwise FDR-corrected *p* = 0.94). We then asked whether vision and language models show a similar convergence as seen in similarity judgements, quantifying the similarity between vision DNN and LLM representations within each dataset (Figure 3b, right). For each model, we selected the layer that best matched the same-modality similarity judgements, and then averaged the RDMs of these layers within each model family for each dataset. Vision DNN and LLM RDMs were moderately correlated in all three datasets (THINGS: Pearson’s *r* = 0.44, NSD: *r* = 0.44, BMD: *r* = 0.42; two-sided stimulus-label permutation tests, all FDR-corrected *p <* 0.001), with no significant differences across datasets (stimulus bootstrap; all pairwise FDR-corrected *p* = 0.86). Vision DNNs and LLMs therefore shared reliable but weaker representational similarity than human vision and language behaviour, and this correspondence was consistent across objects, scenes, and visual events.

After assessing vision-language convergence within behaviour and within models, we further asked how well each model family captured human similarity judgements. Vision DNNs and LLMs explained a significant amount of variance in both vision-based and language-based similarity judgements in every dataset (two-sided Wilcoxon signed-rank tests; FDR-corrected *p <* 0.001; see Figure 3c, top). For each dataset and judgement type, we report the best-performing model and layer within each family (Figure 3c, bottom). The selected vision DNN layers were consistently late-stage representations (final pooling or fully connected layers), whereas the selected LLM representations were all single sentence-level embeddings produced by sentence-embedding models. For static stimuli, vision DNN and LLM model families showed an overall relative advantage over the other in their matching modality. Vision DNNs aligned more strongly than LLMs with vision-based judgements in both THINGS (Pearson’s *r* = 0.17 vs 0.14, FDR-corrected *p <* 0.01) and NSD (*r* = 0.19 vs 0.18; FDR-corrected *p <* 0.05), whereas LLMs aligned more strongly than vision DNNs with language-based judgements in NSD (*r* = 0.15 vs 0.12; FDR-corrected *p <* 0.05), with no difference between model families in THINGS (*r* = 0.093 vs 0.089; *p* = 0.44). For the dynamic video dataset (BMD), LLMs aligned more strongly than vision DNNs with both vision-based (*r* = 0.27 vs 0.17) and language-based (*r* = 0.19 vs 0.07) judgements (FDR-corrected *p <* 0.001). LLM alignment was also higher with vision-based similarity judgements than with language-based similarity judgements (two-sided Mann-Whitney U test; *p <* 0.01). Thus, in BMD, the structure of vision-based similarity judgements was better captured by representations from text-only LLMs than from video-trained vision DNNs.

To examine whether the alignment between models and human similarity judgements remained comparable across stimulus types, we tested for systematic trends from objects to scenes to dynamic events. Across datasets, LLM alignment with vision-based similarity judgements increased across stimulus types (two-sided Jonckheere-Terpstra test; *z* = 4.62, *p <* 0.001), even after controlling for the noise ceiling (*z* = 3.69, *p <* 0.001), with the largest increase from scenes to dynamic events. The increase in LLM alignment with language-based judgements across datasets (*z* = 3.74, *p <* 0.001) did not survive the noise ceiling control (*z* = 1.54, *p* = 0.12). Vision DNN alignment did not change significantly across stimulus types for either vision-based or language-based similarity judgements, before or after controlling for the noise ceiling (two-sided Jonckheere-Terpstra and Kruskal-Wallis tests, all *p >* 0.05). LLMs therefore aligned increasingly well with vision-based similarity judgements from objects to scenes to dynamic events, whereas vision DNN alignment remained at a similar level across stimulus types.

### Vision DNNs align best with early and mid-level visual areas, whereas LLMs and behavioural judgements align best with face– and body-selective regions

Our results showed that vision– and language-based similarity structures converged strongly in behaviour and more weakly in models. To examine whether this correspondence extends to neural representations, we assessed how behavioural and model RDMs aligned with brain RDMs across visual cortex, and whether vision and language predictors explained shared or distinct neural variance. We restricted cross-dataset analyses to functionally defined ROIs available in all three datasets.

Behaviour-brain alignment increased from early and mid-level (V1, V2, V3, and hV4) to higher-level visual cortex across all three datasets, though the relative strength of the two modalities differed by dataset (Figure 4). Both vision and language behavioural RDMs aligned significantly with the brain RDMs in all ten ROIs in every dataset (two-sided Wilcoxon signed-rank tests for NSD and BMD, two-sided stimulus-label permutation tests for THINGS; FDR-corrected *p <* 0.05). Alignment for both modalities was strongest in higher-level visual areas, particularly the face– and body-selective regions, and weakest in early visual areas across all datasets. The relative contributions of the two modalities diverged across higher-level regions. In scene-selective regions, vision behaviour exceeded language behaviour in parahippocampal place area (PPA) in THINGS and BMD, and retrosplenial cortex (RSC) in NSD and BMD. In face– and body-selective regions (fusiform face area (FFA), occipital face area (OFA), extrastriate body area (EBA)), the pattern was dataset-dependent: the two modalities did not differ in THINGS, language mostly exceeded vision in NSD (FFA and EBA), and vision exceeded language in BMD (all FDR-corrected *p <* 0.05). Across datasets, most of the explained brain variance was shared between vision and language behaviour rather than unique to either modality (Figure 4). This shared component averaged 73% of the explained variance in THINGS, 59% in NSD, and 64% in BMD, and significantly exceeded the unique contributions of either modality in higher-level visual ROIs in all datasets. Vision and language behaviour thus predicted largely overlapping brain representations across all stimulus types.

**Fig. 4.**
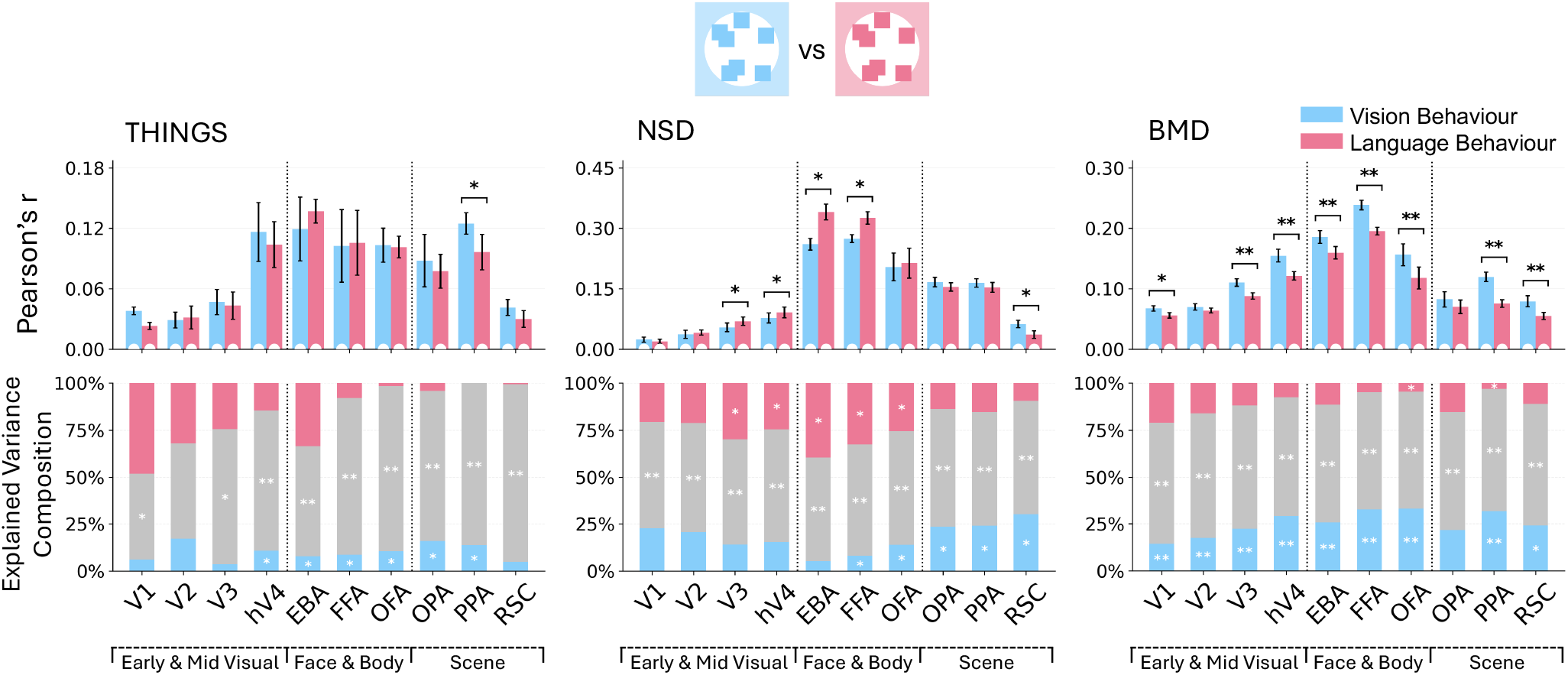
Behaviour-brain representational alignment and variance partitioning. Top: Alignment between behavioural RDMs (vision, language) and brain RDMs across ROIs for each dataset. The alignment was quantified using Pearson’s correlation. White dewdrops at the base of each bar indicate a correlation significantly greater than zero, and brackets with asterisks denote significant differences in the amount of variance that vision and language behaviour explain in each ROI. Both were tested using two-sided Wilcoxon signed-rank tests for NSD and BMD, and stimulus-label permutation tests for THINGS. Alignment relative to the noise ceiling is provided in Supplementary Figure 6. Bottom: Variance explained uniquely by vision behaviour, uniquely by language behaviour, and shared between vision and language behaviour, shown as percentages of total explained variance across ROIs for each dataset. Significance was assessed using the same tests as above, applied within each dataset. All p-values are FDR-corrected across the ten ROIs. Significance: \**p <* 0.05, \*\**p <* 0.01, \*\*\**p <* 0.001.

Unlike behaviour-brain alignment, model-brain alignment did not increase consistently from early to higher-level visual cortex (Figure 5a). Vision DNNs aligned most strongly with early to mid-level visual areas, where they exceeded LLMs in all datasets; in NSD, vision DNNs also exceeded LLMs in all scene-selective regions. LLMs, by contrast, aligned most strongly with face– and body-selective regions, although this peak alignment was least pronounced in THINGS. In face– and body-selective regions, LLMs matched the variance explained by vision DNNs across datasets and explained significantly more variance in FFA and EBA in BMD. Variance partitioning showed that vision DNNs and LLMs had a smaller proportion of the shared explained variance (Figure 5a), as compared to vision and language behaviour (Figure 4). In early to mid-level visual areas, vision DNNs explained more unique variance than LLMs, and vision DNNs’ unique contribution accounted for most of the explained variance across datasets (THINGS: 86%; NSD: 87%; BMD: 69%). In all higher-level visual regions, the two model families shared a significant portion of the explained variance (FDR-corrected *p <* 0.05), but this shared component was smaller than that between vision and language behaviour (mean across ROIs, models vs behaviour; THINGS: 54% (models) vs 82% (behaviour); NSD: 28% (models) vs 60% (behaviour); BMD: 40% (models) vs 64% (behaviour)). Vision DNNs and LLMs thus each explained significant unique variance in nearly all higher-level regions across datasets.

**Fig. 5.**
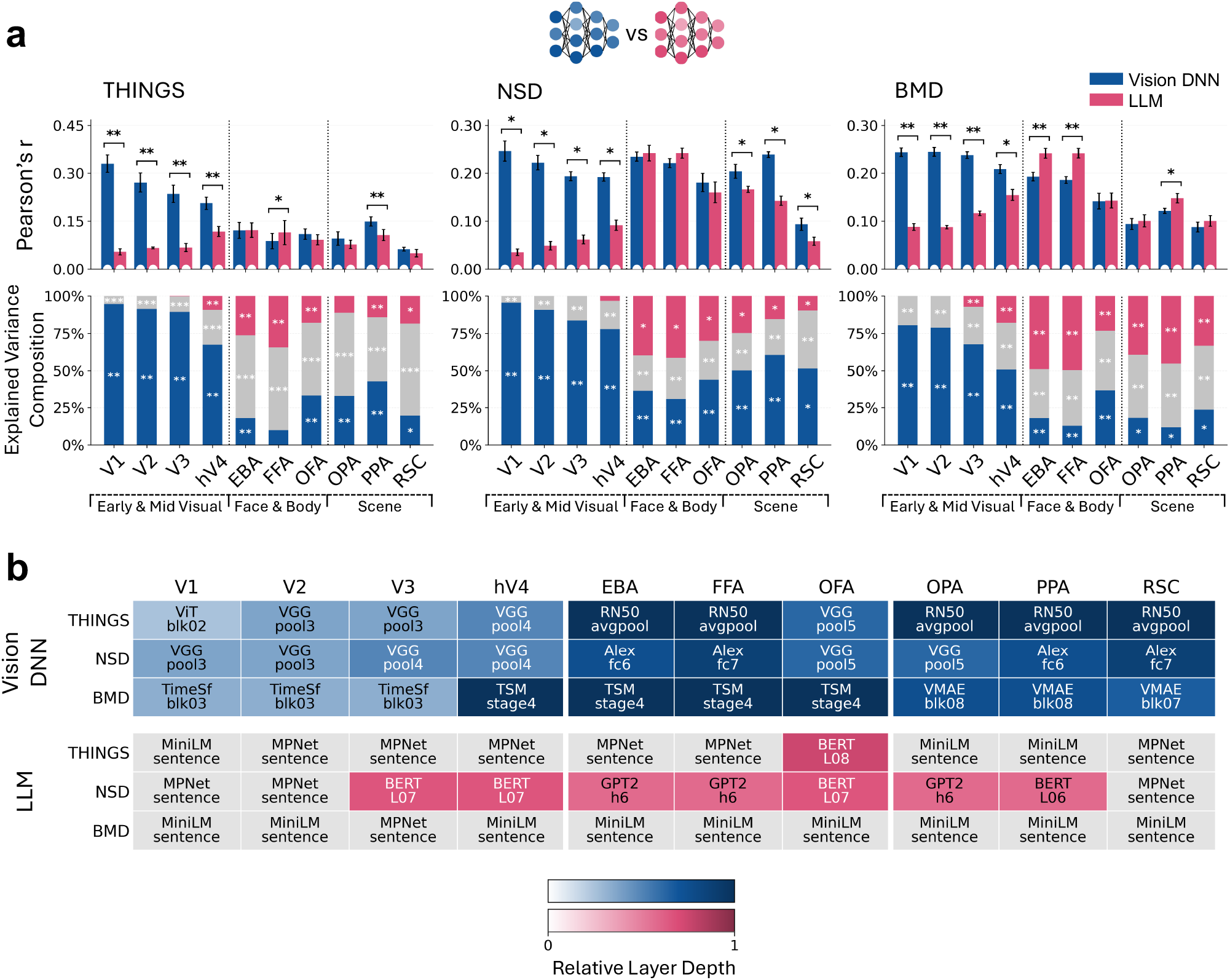
Brain alignment and variance partitioning of vision DNNs and LLMs, and the hierarchy of the best-matching model layers across ROIs. *a*, Model-brain representational alignment and variance partitioning. Top: Alignment between model RDMs (vision DNN, LLM) and brain RDMs across ROIs for each dataset. Alignment relative to the noise ceiling is provided in Supplementary Figure 6. Bottom: Variance explained uniquely by vision DNN, LLM, and shared between vision DNN and LLM, shown as percentages of total explained variance across ROIs for each dataset. Conventions are the same as in Figure 4. *b*, Depth of the best-matching vision DNN layer across ROIs. Top three blocks (THINGS, NSD and BMD, from top to bottom): For each vision DNN and ROI, the best-matching layer was selected by cross-validation. We used image DNNs for THINGS and NSD, and video DNNs for BMD. Bottom block: Summary of the selected LLM models and layers across ROIs for each dataset. Shading indicates relative layer depth, calculated as the layer position divided by the total number of extracted layers in each model (1 = final layer; SlowFast’s slow and fast pathways are counted separately).

The alignment values described above reflect the best-matching model and layer for each ROI, selected by cross-validation separately within each model family. We examined which models and layers were selected, and whether layer depth tracked the visual hierarchy (Figure 5b). For vision DNNs, the relative depth of the selected layers generally increased along the visual hierarchy across all three datasets, with intermediate layers best matching early to mid-level visual areas and later layers best matching higher-level visual areas. For static stimuli, DNN-based image models were selected for nearly all ROIs. VGG16 was selected for V1–hV4 and OFA (except V1 in THINGS, for which the vision transformer ViT-B/16 was selected), whereas ResNet50 (THINGS) and AlexNet (NSD) were selected for EBA, FFA, PPA and RSC. OPA was best matched by ResNet50 in THINGS and by VGG16 in NSD. For dynamic events, the video transformer TimeSformer was selected for V1–V3, the DNN-based video model TSM-R50 for hV4 and the face– and body-selective regions (EBA, FFA, OFA), and the video transformer VideoMAE-B for the scene-selective regions (OPA, PPA, RSC). For LLMs, sentence embedding models (MPNet and MiniLM) were selected in nearly all ROIs in THINGS and BMD (except for occipital face area (OFA) in THINGS). In NSD, MPNet and MiniLM were selected only in V1, V2 and RSC, whereas intermediate layers of BERT or GPT-2 best matched the remaining ROIs. The vision DNN results are consistent with previous reports of hierarchical layer-to-region correspondence (Cichy et al., 2016; Güçlü and van Gerven, 2015). Similarly, the LLM results are consistent with previous findings that sentence embeddings of image captions align with higher-level visual cortex (Doerig et al., 2025) and that caption representations from contextual language models best predict high-level visual cortex at intermediate network depth (Bavaresco et al., 2026).

### A cross-modal shared component dominates the explained variance across higher-level visual cortex

To test whether predictors that differed in modality (vision or language) and system (human judgements or computational models) accounted for overlapping or distinct components of brain variance, we fitted all four predictors jointly and partitioned the explained variance into the unique contribution of each predictor and the variance shared among predictors (Figure 6). Because these variance-partitioning components are computed as differences between cross-validated *R*^2^ values, they can be negative. We retained negative values for statistical testing but set them to zero for visualisation and reported percentages. On average, negative components were small (mean 2.5% across ROIs) and arose primarily from the unique contributions of the behavioural predictors.

**Fig. 6.**
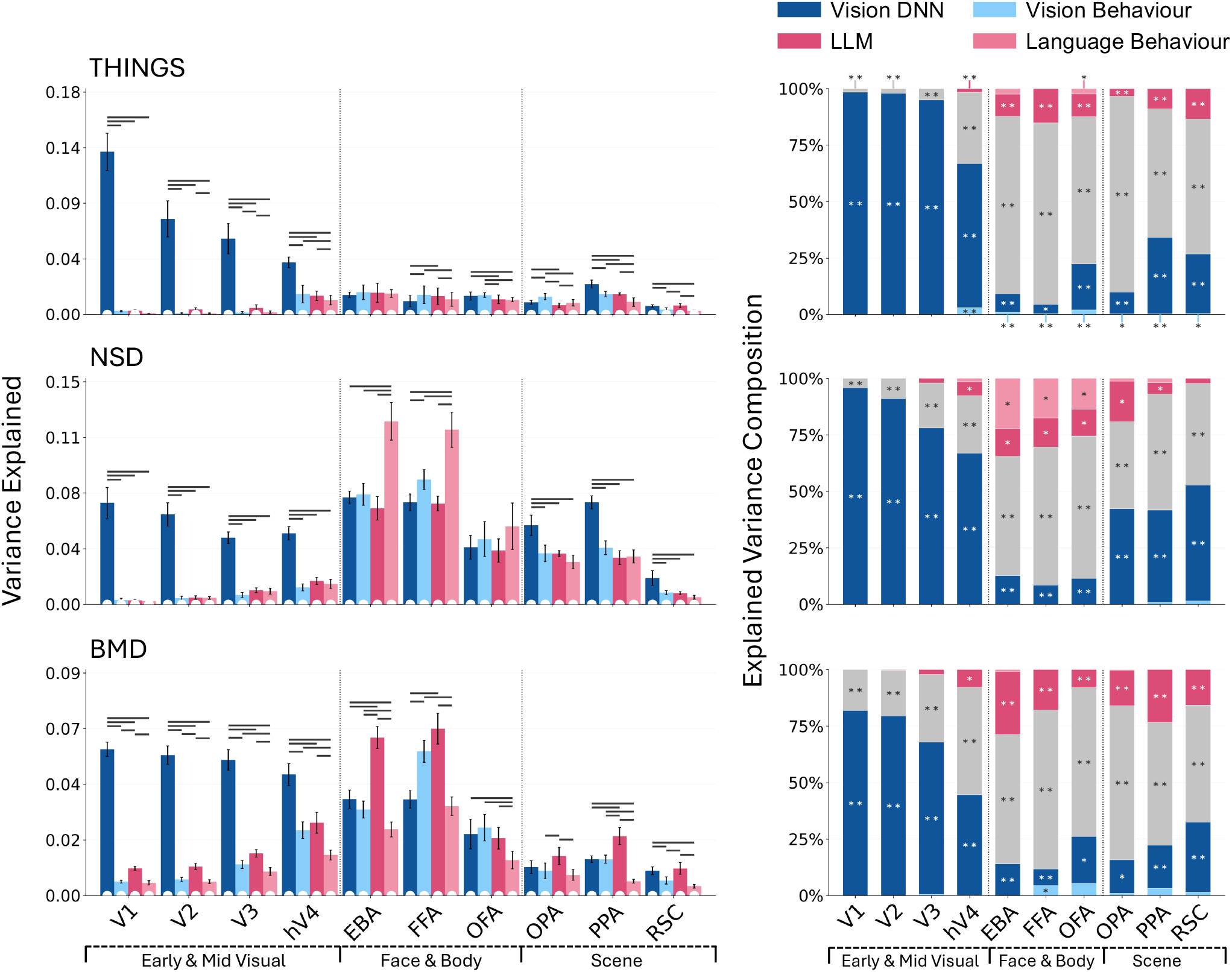
Variance in brain RDMs explained by vision and language predictors from human behaviour and from models, and the variance partitioning. For each ROI and participant, four predictor RDMs were entered into a single non-negative least-squares fit: group-level vision-based similarity judgements (vision behaviour) and language-based similarity judgements (language behaviour), and the best-fitting layer of vision DNNs and of LLMs, each selected per ROI using cross-validation. Variance explained was quantified as the mean across splits of the squared held-out Pearson’s correlation, with negative correlations set to zero. Left: Variance explained by each predictor fitted alone, across ROIs for each dataset. Because variance explained by the predictors overlap heavily, these bars are not additive and sum to more than the joint fit. White dewdrops at the base of each bar indicate variance explained significantly greater than zero. Dark grey horizontal lines above the bars join pairs of predictors that differ significantly in the amount of variance they explain when fitted alone. Right: Percentage of total explained variance attributable uniquely to each of the four predictors, and shared between two or more of them. Negative unique contributions (which cross-validation can produce, see Methods) are set to zero for display but tested unclipped. Significance was assessed with two-sided Wilcoxon signed-rank tests for NSD and BMD, and participant-level two-sided stimulus-label permutation tests for THINGS. All *p*-values are FDR-corrected across the ten ROIs. Significance: \**p <* 0.05, \*\**p <* 0.01, \*\*\**p <* 0.001.

In early to mid-level visual areas, vision DNNs accounted for most of the variance explained by the joint four-predictor model (THINGS: 86%, NSD: 82%, BMD: 67%; FDR-corrected *p <* 0.01), consistent with the model-brain alignment results. The component shared across all four predictors was significant in every early to mid-level visual ROI (FDR-corrected *p ≤* 0.01) and the shared component’s contribution increased along the visual hierarchy, from a small fraction of the explained variance in V1 to a substantial proportion of the explained variance in hV4 (THINGS: 3% (V1) to 33% (hV4); NSD: 6% (V1) to 26% (hV4); BMD: 20% (V1) to 48% (hV4)).

In higher-level visual regions, the shared component was significant in all ROIs (FDR-corrected *p <* 0.01), accounting on average for 75% of the explained variance in THINGS, 52% in NSD, and 62% in BMD. This shared variance was predominantly cross-modal. To characterise its composition, we grouped shared terms into those restricted to a single modality (vision DNNs with vision behaviour; LLMs with language behaviour) and those spanning both modalities. On average, 91% of the shared component (ranging 76–99%) was cross-modal, and this cross-modal component was itself significant in almost every higher-level ROI (FDR-corrected *p <* 0.01). Despite this dominance of shared variance, reliable predictor-specific contributions remained. The unique variance of both vision DNNs and LLMs was significant in every higher-level ROI across all three datasets (FDR-corrected *p <* 0.05). Vision behaviour contributed negligibly throughout, and language behaviour contributed unique variance mainly in NSD. LLMs explained more unique variance in face– and body-selective regions than in scene-selective regions in all three datasets (FDR-corrected *p <* 0.01), exceeding all other predictors in EBA and FFA in THINGS and BMD. In NSD, language behaviour explained more variance than all other predictors in EBA and FFA (FDR-corrected *p <* 0.05). Language behaviour also explained most unique variance in both regions, significantly outperforming LLMs in EBA (FDR-corrected *p <* 0.05) and numerically in FFA (FDR-corrected *p* = 0.078). In OFA, language behaviour and LLMs did not differ significantly in either total or unique variance explained. Vision– and language-derived predictors converged on a predominantly shared, cross-modal component of brain activity variance in higher-level visual cortex, while language-derived predictors made the largest unique contributions in EBA and FFA.

The small unique contributions of the behavioural predictors in the four-way partition do not necessarily mean that human judgements add little beyond models. Because vision– and language-based judgements were strongly correlated (*r ≈* 0.6), each absorbed much of the other’s unique contribution. We therefore compared each judgement type with the model of the same modality (Figure 7).

**Fig. 7.**
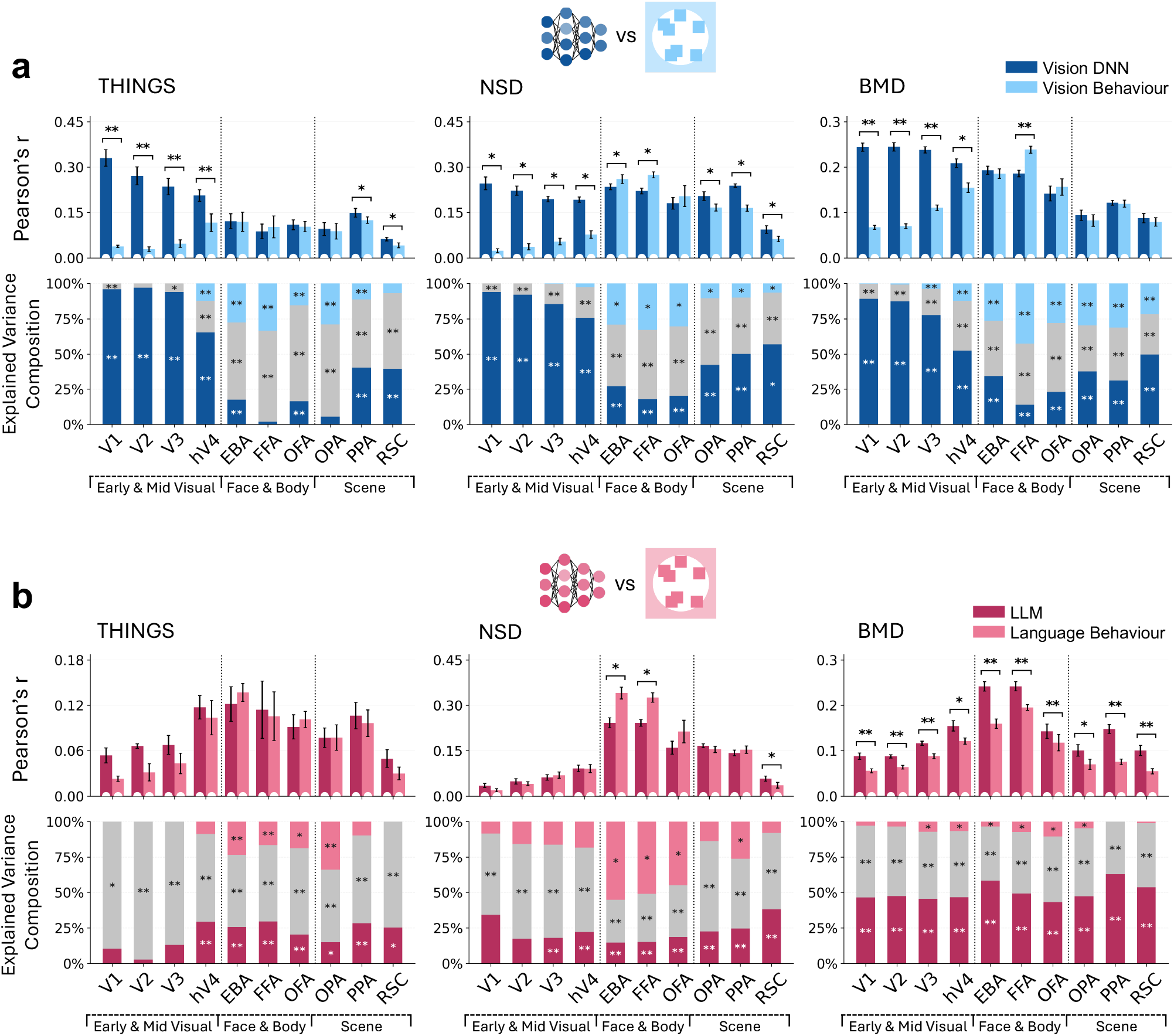
Behaviour-brain and model-brain representational alignment and variance partitioning within each modality. ***a***, Vision DNNs and vision-based similarity judgements. ***b***, LLMs and language-based similarity judgements. Top row of each panel: Alignment of model RDMs and behavioural RDMs with brain RDMs across ROIs for each dataset. Bottom row of each panel: Variance explained uniquely by the model, uniquely by behaviour, and shared between the model and behaviour, as a percentage of total explained variance across ROIs for each dataset. Conventions are the same as in Figure 4.

In early and mid-level visual areas, vision DNNs dominated the vision partition, accounting for most of the explained variance (THINGS: 88%, NSD: 87%, BMD: 77%). Vision-based similarity judgements shared relatively little variance with vision DNNs (9–19%) and contributed little unique variance (less than 4% on average) in early and mid-level visual areas. In contrast, the language partition was dominated by variance shared between LLMs and language-based similarity judgements (THINGS: 84%, NSD: 62%, BMD: 49%), and the unique contribution of language-based judgements was small (2–15%) in early and mid-level visual areas and significant only in V3 and hV4 in BMD. In face– and body-selective regions, behaviour explained significant unique variance in every region and dataset, in both the vision (25–32%) and language (7–50%) partitions. Behaviour’s unique contribution exceeded that of the corresponding model in several regions: vision-based judgements exceeded vision DNNs in EBA and FFA in THINGS, all three regions (FFA, OFA, EBA) in NSD, and FFA and OFA in BMD; language-based judgements exceeded LLMs in all three regions in NSD. Consistent with this result, behaviour also aligned more strongly with the brain than the corresponding model in EBA and FFA in NSD (both modalities) and in FFA in BMD (vision only). In scene-selective regions, behaviour contributed less unique variance than in face– and body-selective regions in every dataset and modality, and the unique model contributions were correspondingly larger in all cases except the language partition in THINGS.

Collectively, our findings indicate that vision– and language-derived predictors overlapped with each other, while retaining unique contributions in explaining brain responses. Vision-language convergence was higher within behaviour than within models, with no difference across datasets. From objects to dynamic events, LLM alignment with vision-based similarity judgements increased, whereas vision DNN alignment remained stable. In early to mid-level visual areas, vision DNN accounted for most of the explained brain variance, with the shared component between predictors growing from V1 to hV4. In higher-level visual cortex, the shared component accounted for the largest share of explained variance, while predictor-specific contributions remained. In particular, the advantage of language-derived predictors in EBA and FFA suggests that these regions may represent semantic information that is not fully captured by vision-derived predictors. Comparing predictors within each modality further revealed that, in face– and body-selective regions, human similarity judgements explained additional variance beyond both vision DNNs and LLMs. Because neither judgement type contributed much unique variance in the four-way partition, this additional unique variance contributed by behaviour within each modality was largely shared between vision– and language-based similarity judgements.

## Discussion

We investigated how vision– and language-derived predictors converge in explaining the representational space of visual cortex. Building on previous evidence of a shared contribution of vision and language in predicting patterns of brain activity, we asked how this convergence holds across predictor types (human behaviour vs models) and stimulus types (objects, scenes, and dynamic events), and how the vision-language convergence varies across the visual hierarchy and heterogeneity of visual brain areas. Our findings reveal substantial but incomplete vision-language convergence and clarify the shared and unique contributions of vision– and language-based representations across stimulus types and brain regions.

Across stimulus types, the shared vision and language component accounted for more than half of the explained variance in higher-level visual regions, and its largest term was common to all four predictors. Because this shared component increased along the visual hierarchy, vision-language convergence appears to occur primarily at a more abstract and conceptual level, in line with the progressive emergence of semantic information along the ventral and lateral visual pathways (Devereux et al., 2018; Wurm and Caramazza, 2022). Previous work also showed that vision– and language-based similarity judgements of natural scenes explain largely shared variance in brain responses of bilateral mid-to high-level visual areas, with only modest modality-specific contributions (Simkova et al., 2026). Our results extend this finding to objects and dynamic events, and from human similarity judgements to vision DNNs and LLMs, separating the unique and shared contributions across all four predictors. At the level of functionally-defined ROIs, we further found that the vision-language convergence increases along the visual hierarchy and varies across category-selective regions. This convergence is compatible with the Platonic Representation Hypothesis (PRH), which proposes that artificial neural networks trained on different modalities increasingly converge on a shared representation of the world (Huh et al., 2024). Yet vision DNNs and LLMs also explained unique variance in higher-level visual regions, with a similar pattern in all three datasets: LLM-specific variance was greatest in face– and body-selective regions, and vision-DNN-specific variance was greatest in scene-selective regions.

These unique contributions suggest that convergence between vision and language is partial, which may reflect modality-specific constraints. Recent studies have found weaker and more limited cross-modal convergence than originally reported, indicating that each modality retains distinct information (Koepke et al., 2026; Gröger et al., 2026; Tjandrasuwita et al., 2025). Umwelt Representation Hypothesis (URH; Bosch et al., 2026) proposes that each system learns representations shaped by its own ecological constraints. These constraints include the environment, sensory and motor capacities and goals for humans; and the training data, architecture and learning objectives for artificial networks. Representations of different systems therefore align to the extent that their constraints overlap and tend to diverge where these constraints differ. Differences between systems thus would reflect adaptation to distinct constraints. This framework may also explain why vision-language convergence was stronger in human judgements (*r ≈* 0.6) than between models (*r ≈* 0.4), as humans acquire visual and language knowledge under shared ecological constraints, whereas vision DNNs and LLMs are trained on distinct inputs and objectives.

Vision DNNs and LLMs made distinct contributions across the visual cortex. In early and mid-level visual areas, vision DNNs accounted for most of the explained variance, whereas LLMs and behavioural predictors showed weaker brain alignment and explained little unique variance. This pattern is consistent with the sensitivity of these areas to low-level visual features, which are well captured by the early and intermediate layers of vision DNNs (Yamins et al., 2014; Güçlü and van Gerven, 2015; Yamins and DiCarlo, 2016). In face– and body-selective regions, LLMs explained significant unique variance across all three datasets, and their brain alignment matched or exceeded that of vision DNNs, particularly in EBA and FFA. Beyond their preferred categories, both regions encode a range of person-related and social information. EBA represents visual social cues such as body posture (Marrazzo et al., 2023), interpersonal distance and facingness (McMahon et al., 2023), and also distinguishes different types of social interaction (Walbrin and Koldewyn, 2019). Similarly, FFA represents social attributes beyond face identity, including perceived social traits and gender (Tsantani et al., 2021). More broadly, representations in lateral occipitotemporal cortex are organised by the social content and the purpose of observed actions (Wurm et al., 2017; Tarhan and Konkle, 2020), and these representations extend beyond the canonical action observation network into ventral temporal cortex, including the fusiform gyrus (Han et al., 2026). LLM embeddings appear to capture these properties as well. LLMs explain variance in human similarity judgements of action videos and their descriptions beyond hand-labelled semantic features, mainly through action-related information (Dima et al., 2024), and predict human ratings of actions and social interactions better than vision models (Garcia et al., 2025). This sensitivity to action-related and social information may help explain the unique variance LLMs contribute in EBA and FFA. In scene-selective regions, vision DNNs explained more unique variance than LLMs for static stimuli (THINGS and NSD), whereas the two model families made comparable contributions for dynamic events (BMD; see Figure 6). This difference may reflect the greater relevance of action and contextual information in dynamic events, which static images tend to underrepresent (Haxby et al., 2020).

Our findings extend previous evidence that language-derived representations align with high-level visual cortex (Doerig et al., 2025; Wang et al., 2023; Simkova et al., 2026) by identifying where LLMs explain unique variance beyond vision DNNs and how these contributions vary across stimulus types. Compared with humans, vision DNNs place greater weight on visual than semantic dimensions (Mahner et al., 2025), whereas LLMs capture conceptual and relational information, with perceptual knowledge limited by what can be learned from text (Lewis et al., 2019; Marjieh et al., 2024; Wang et al., 2026). The specific dimensions underlying the variance uniquely explained by LLMs remain to be identified, for example through interpretable feature-based models of objects, actions and social interactions (Contier et al., 2024; McMahon et al., 2023). Future work could also examine other brain regions, such as superior temporal sulcus, which is involved in social perception (Pitcher and Ungerleider, 2021), and anterior temporal lobe, which has been proposed to serve as a transmodal semantic hub (Lambon Ralph et al., 2017).

We found strong convergence between human similarity judgements of visual stimuli and of their language descriptions, consistent with previous findings for action videos and scenes (Dima et al., 2024; Simkova et al., 2026). Because participants received no explicit instructions about which properties to consider in the multi-arrangement task, this convergence suggests that the dimensions people spontaneously use to judge similarity are largely expressible in language, in line with evidence that many dimensions underlying object similarity judgements are interpretable and nameable (Hebart et al., 2020; Jozwik et al., 2022, 2023). We further showed that vision-language convergence was comparable across objects, scenes, and dynamic events, even though these stimulus types differ in the kinds and amounts of information they contain. This finding suggests that people judge similarity at a similar level of abstraction across stimulus types, relying also on the overall meaning of the stimulus, which brief text descriptions can convey. In the visual cortex, both vision– and language-based similarity judgements aligned most strongly with face– and body-selective regions, and language-based judgements generally explained little unique variance beyond vision-based judgements (Figure 4). An exception emerged in NSD, where language-based judgements contributed unique variance in face– and body-selective regions, consistent with findings by Simkova et al. (2026). Given that the behavioural judgement tasks were identical across datasets, this difference may relate to the tasks in fMRI experiments. NSD used a continuous recognition memory task (Allen et al., 2022), which may encourage the encoding of verbally expressible information, whereas THINGS and BMD used catch-trial detection (Hebart et al., 2023; Lahner et al., 2024), although differences in stimuli cannot be ruled out. Together, vision– and language-based similarity judgements explained variance beyond that captured by computational models (Figure 7). Much of this variance was shared between vision– and language-based judgements (Figure 6) and concentrated in face– and body-selective regions, in contrast to the models that explained unique variance throughout visual cortex (see also Supplementary Figure 7). Human similarity judgements may thus emphasise socially meaningful dimensions accessible through both seeing and reading, while models also capture finer-grained visual and language-based features that people do not explicitly use when judging similarity (Contier et al., 2024; Groen et al., 2018). Human behaviour and models may therefore provide complementary information about high-level visual representations, highlighting the value of incorporating human behaviour into model evaluation and training (Muttenthaler et al., 2025; Sucholutsky et al., 2025).

In summary, vision-language convergence was higher within behaviour than within models. The shared representational space captured by vision– and language-derived predictors grew along the visual hierarchy. LLMs and shared representations of vision– and language-based human similarity judgements explained variance not captured by vision DNNs, most clearly in face– and body-selective regions (FFA and EBA). We find a partially shared representational space for vision and language in high-level visual cortex, most clearly in regions representing social information related to people and their actions. As each predictor retained unique variance, combining computational models and human behaviour across vision and language may offer a more comprehensive account of high-level visual representations.

## AUTHOR CONTRIBUTIONS

QZ and KMJ designed the experiments. QZ performed the analyses. QZ and KMJ wrote the paper. KMJ supervised the work.

## DATA AVAILABILITY

The datasets generated during the current study are available from the corresponding author upon request.

## CODE AVAILABILITY

The code generated during the current study is available from the corresponding author upon request.

## ACKNOWLEDGEMENTS

We thank Annabel Man for coordinating behavioural data collection. We are grateful to Jacek Podlaski and Timothée Maniquet for critically reading the manuscript and assisting with proofreading. This research was funded in part by the Wellcome Trust [206521/Z/17/Z] and the Royal Society [URF\R1\251359]. For the purpose of open access, the author has applied a CC BY public copyright licence to any Author Accepted Manuscript version arising from this submission.

## USE OF GENERATIVE AI

GPT-4.1 mini (OpenAI) synthesised representative captions for each stimulus from its five human annotations (see Methods). We reviewed every generated caption. Claude Opus 5 (Anthropic) assisted with debugging and with the analysis and figure code. Claude Opus 5.5 (Anthropic) and Grammarly (v.1.2.306.1984) suggested improvements to the manuscript structure and helped polish the language. The authors directed, reviewed, and verified all AI-assisted code and text, checked every reported value against the data and every reference against its source, and take full responsibility for the content of this manuscript. The study design, analyses, interpretation of the results, and the wording of the manuscript are the authors’ own.

