## Supplementary_Information for "Partially shared representational space for vision and language across objects, scenes, and visual events"

### Stimuli and caption generation

#### 1. THINGS Labelling Task

We collected human-written captions for 100 images from the THINGS dataset, following the captioning instructions used in Microsoft COCO (Chen et al., 2015). Five participants were recruited via Prolific (www.prolific.com), and each annotated the full set of 100 images, yielding 500 captions in total (five per image). Participants received the following instructions.

> Please provide a one-sentence caption for each picture (8–15 words), and follow these instructions:
>
> - Describe all the important parts of the picture.
> - Do not start the sentences with "There is".
> - Do not describe unimportant details.
> - Do not describe things that might have happened in the future or past.
> - Do not describe what a person might say.
> - Do not give people proper names.
> - The sentences should contain 8–15 words.

#### 2. Caption Aggregation Prompts for GPT

**Static Images:**
You are given five annotations describing the same image. The task is to generate one sentence to describe that image based on the converging information found in the five annotations.

**Dynamic Video Clips:**
You are given five annotations describing the same 3-second video clip of daily scenes. The task is to generate one sentence to describe that video clip based on the converging information found in the five annotations.

**Requirements:**

- The sentence should contain at least eight words, but should not exceed the word count of the longest annotation.
- If a detail is mentioned in multiple annotations, include that detail in your response.
- Do not include a specific detail if it appears in only one annotation.
- If annotations contain conflicting details, try to use more general descriptions to accommodate multiple possibilities.
- Keep the wording and level of detail close to the original annotations.
- Use simple, literal image-description language.
- Write one sentence only.
- Avoid redundant or uninformative phrases.
- The sentence should be grammatically correct and natural.

### Supplementary Results

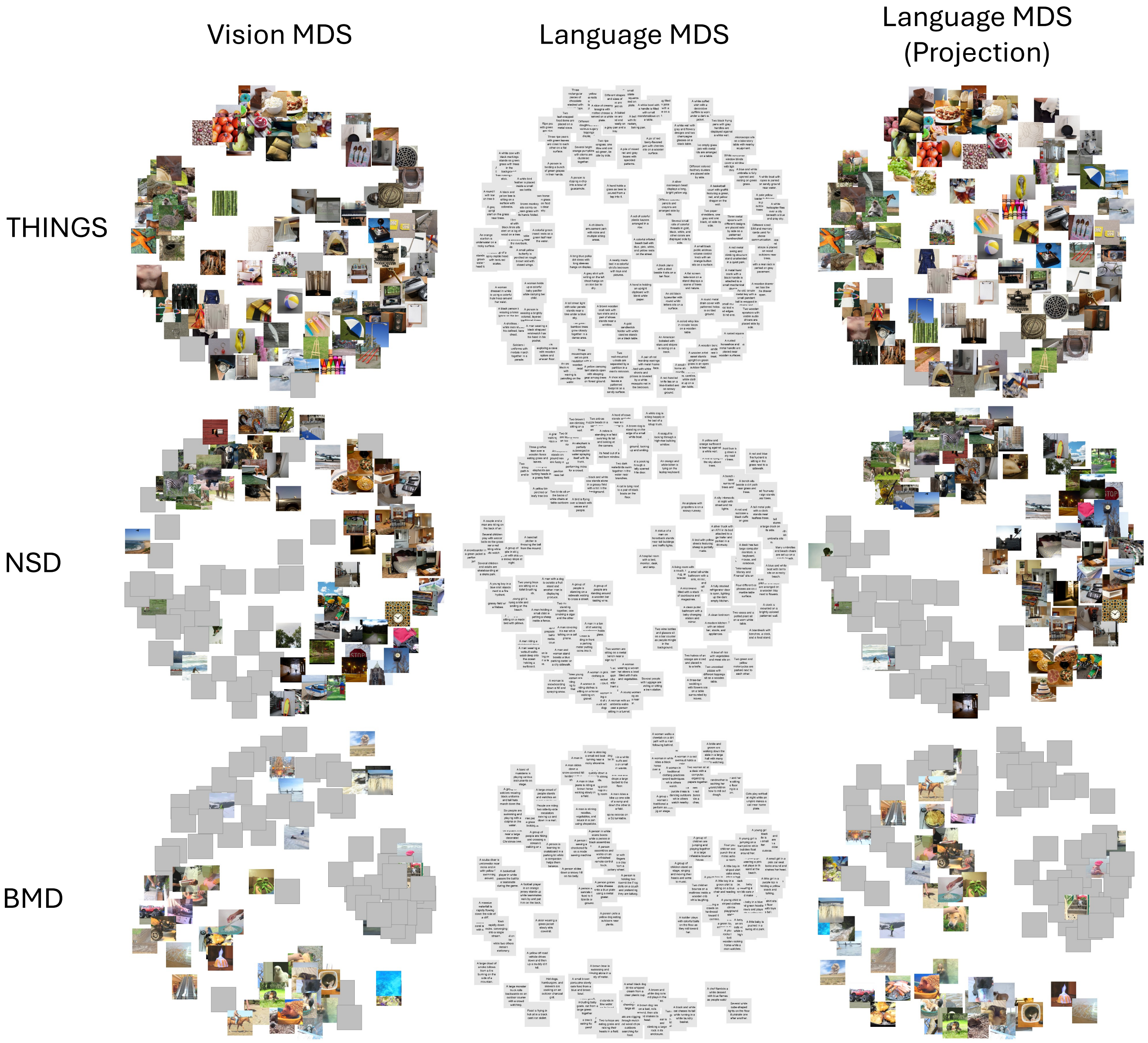


**Supplementary Figure 1. Multidimensional scaling (MDS) of behavioural similarity judgements for vision and language.** Each panel shows a two-dimensional metric MDS embedding of the group-averaged RDM from the MA tasks, for THINGS (top row; 100 stimuli), NSD (middle row; 100 stimuli) and BMD (bottom row; 102 stimuli). Stimuli judged as more similar are placed closer together, and stimuli judged as less similar are placed further apart. Vision MDS (left): similarity structure based on participants' arrangements of the visual stimuli. Grey squares are placed over face-containing images in line with bioRxiv policy. Language MDS (middle): similarity structure based on participants' arrangements of the language descriptions. Language MDS (Projection) (right): the same language similarity structure, with each caption replaced by the image or video it describes. This column is a visual aid for comparing the language and vision arrangements directly. Participants did not see any images or videos in the language task. BMD videos are shown as their middle frame here.

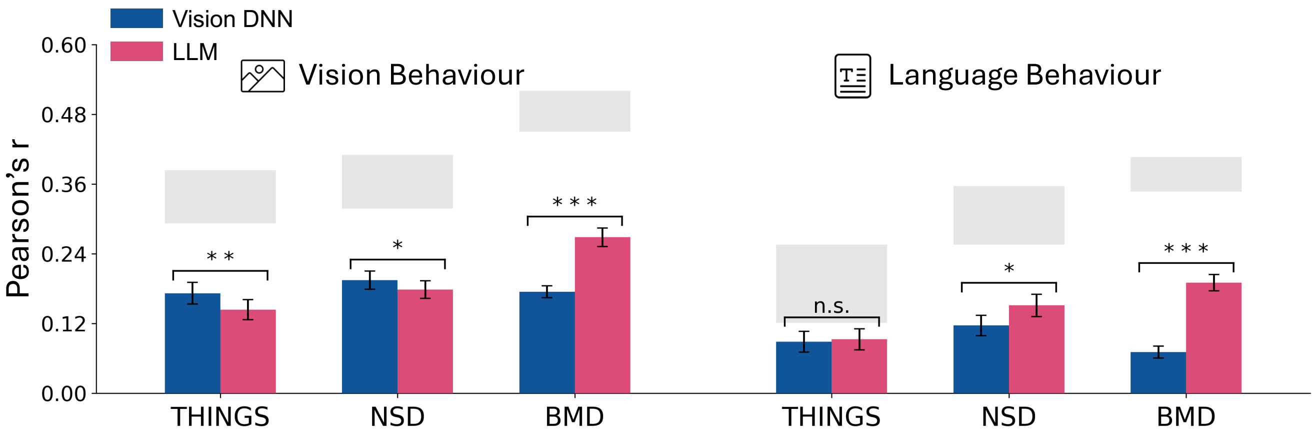


**Supplementary Figure 2. Model-behaviour alignment relative to the behavioural noise ceiling.** Conventions are the same as in Figure 3c. Light grey bars indicate the noise ceiling (Nili et al., 2014), which estimates the range of achievable alignment given inter-participant variability. The lower bound is the mean leave-one-out correlation between each participant's RDM and the average RDM of the other participants. The upper bound is the mean correlation between each participant's RDM and the group-mean RDM (including that participant).

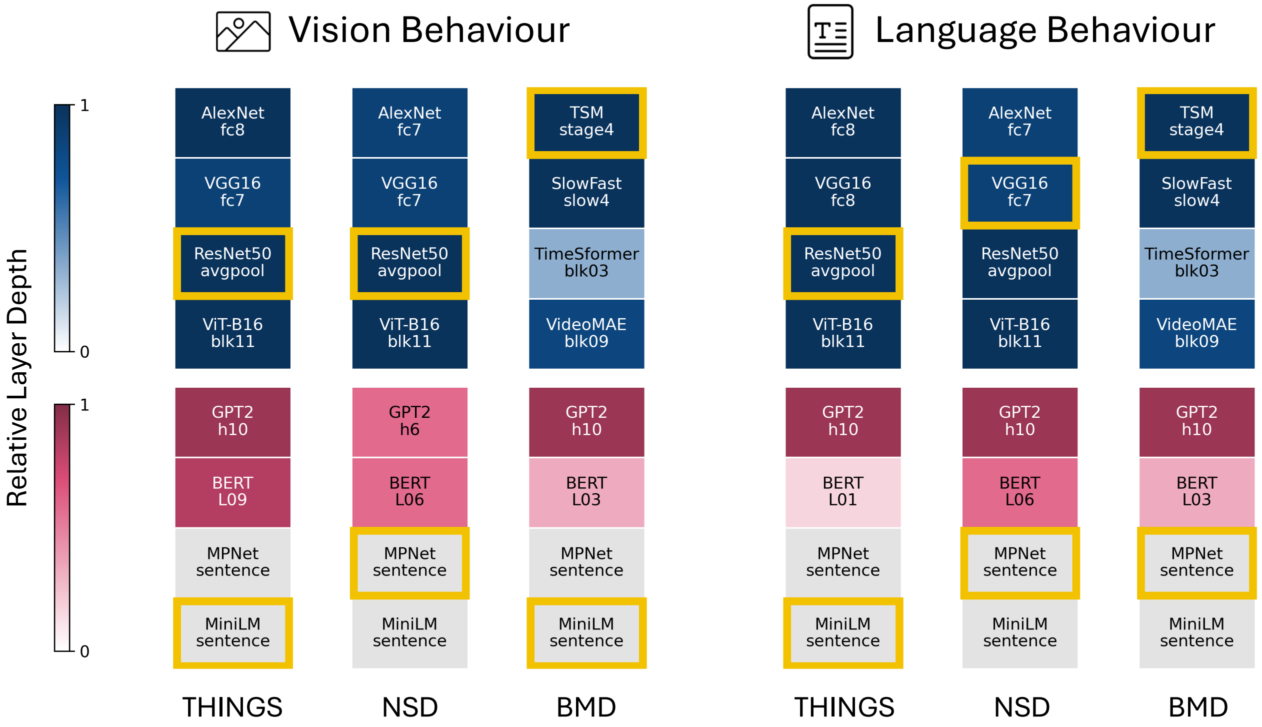


**Supplementary Figure 3. Best-matching model layers for vision- and language-based similarity judgements.** Each column corresponds to one dataset and judgement task, aligned with the bars in Figure 3c in the main text (left three: vision-based judgements, right three: language-based judgements). For each model, the best-matching layer was selected by leave-one-participant-out cross-validation. The upper four rows show vision DNNs and the lower four rows show LLMs. Yellow boxes mark the model selected across each family, which is the one used for the alignment analysis in Figure 3c. Shading indicates relative layer depth, the layer position divided by the number of extracted layers in that model (1 = final layer). MPNet and MiniLM provide a single sentence embedding and are shown in grey.

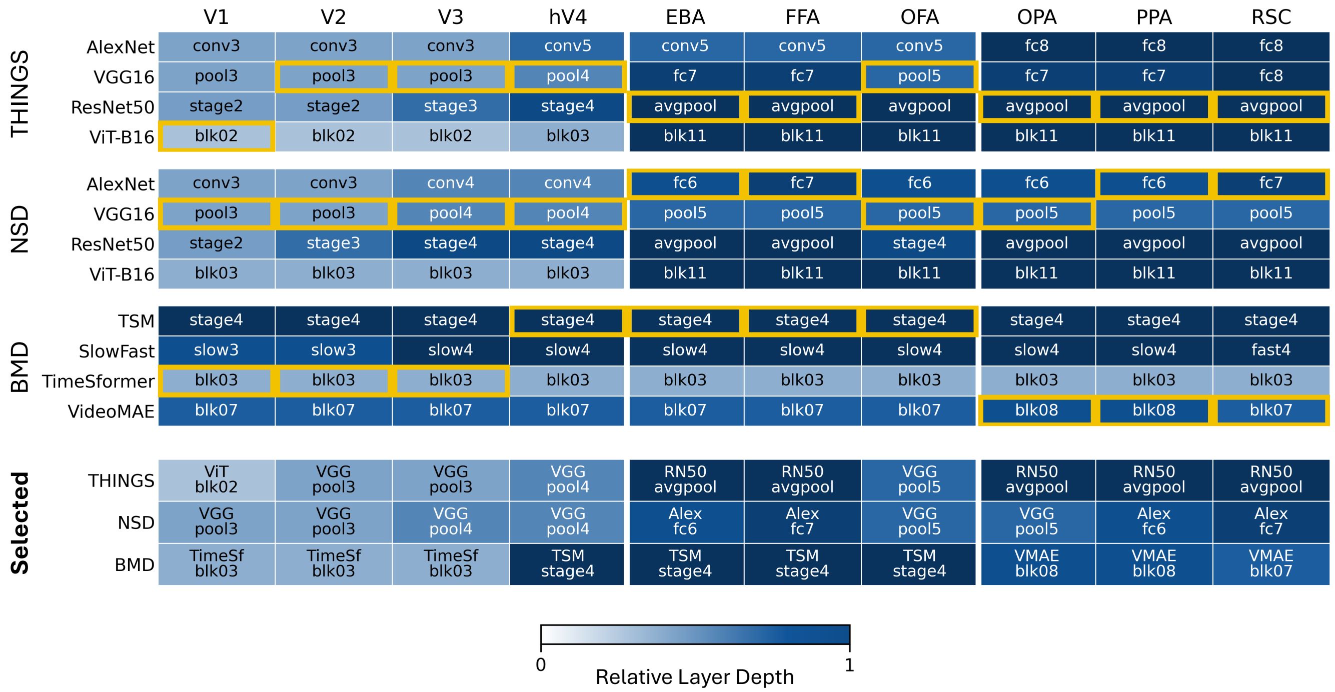


**Supplementary Figure 4. Depth of the best-matching vision DNN layer across ROIs.** Top three blocks (THINGS, NSD and BMD, from top to bottom): For each vision DNN and ROI, the best-matching layer was selected by cross-validation. Yellow boxes indicate the model and layer selected across all vision DNNs for each ROI. Bottom block: Summary of the selected models and layers across ROIs for each dataset. Shading indicates relative layer depth, calculated as the layer position divided by the total number of extracted layers in each model (1 = final layer; SlowFast's slow and fast pathways are counted separately). THINGS and NSD use image DNNs; BMD uses video DNNs.

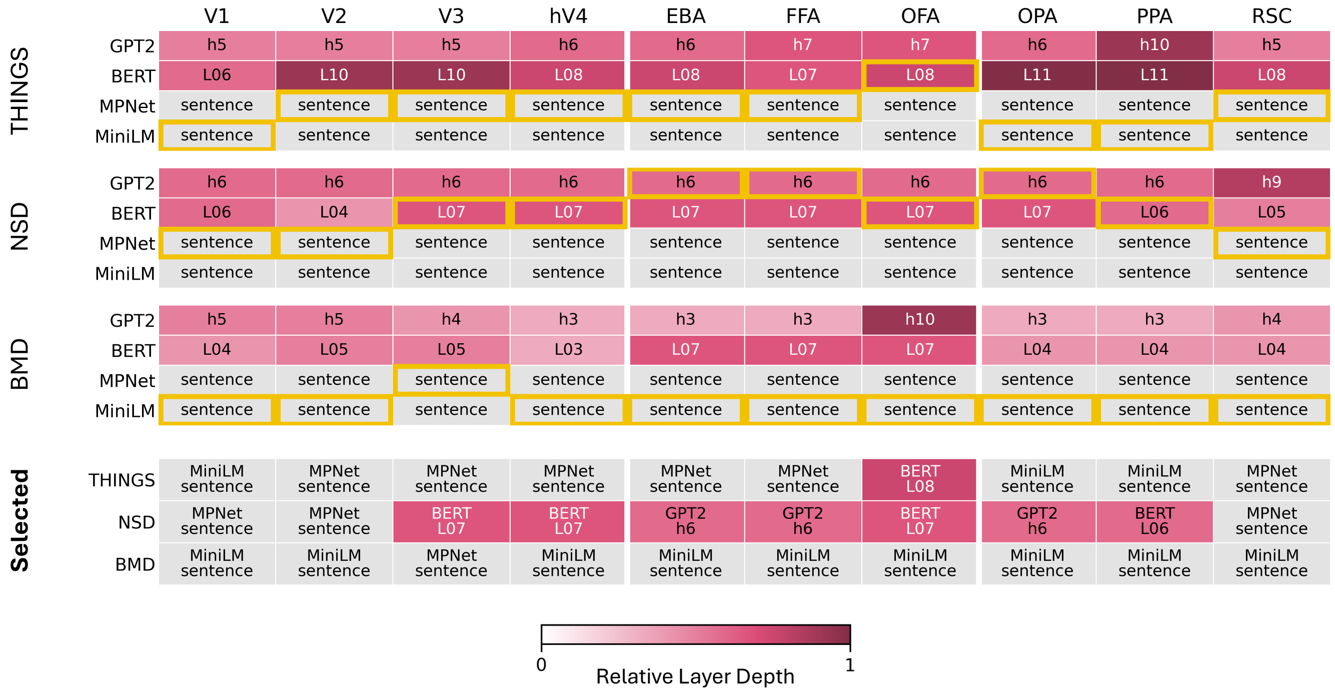


**Supplementary Figure 5. Depth of the best-matching LLM layer across ROIs.** Top three blocks (THINGS, NSD and BMD, from top to bottom): For each LLM and ROI, the best-matching layer was selected by cross-validation. Yellow boxes indicate the model and layer selected across all LLMs for each ROI. Bottom block: the selected LLM and layer for each ROI and dataset. Shading indicates relative layer depth: the layer's position divided by the number of extracted layers in that model (1 = final layer). MPNet and MiniLM provide a single sentence-level embedding and therefore have no layer depth; they are shown in grey.

**Supplementary Table 1. Best vision DNN and LLM layers selected for each ROI: THINGS (objects).** Stability is the proportion of the 10 cross-validation splits in which the reported layer was selected.

| ROI | Vision DNN | Layer | Position | Depth | Stability | LLM | Layer | Position | Depth | Stability |
|---|---|---|---|---|---|---|---|---|---|---|
| V1 | ViT-B16 | blk02 | 3/12 | early | 0.7 | MiniLM | sentence | N/A | N/A | 0.5 |
| V2 | VGG16 | pool3 | 3/8 | mid | 0.8 | MPNet | sentence | N/A | N/A | 0.7 |
| V3 | VGG16 | pool3 | 3/8 | mid | 0.7 | MPNet | sentence | N/A | N/A | 0.5 |
| hV4 | VGG16 | pool4 | 4/8 | mid | 0.7 | MPNet | sentence | N/A | N/A | 0.7 |
| EBA | ResNet50 | avgpool | 5/5 | late (final) | 0.9 | MPNet | sentence | N/A | N/A | 0.9 |
| FFA | ResNet50 | avgpool | 5/5 | late (final) | 0.9 | MPNet | sentence | N/A | N/A | 0.8 |
| OFA | VGG16 | pool5 | 5/8 | mid | 0.6 | BERT | L08 | 9/12 | late | 0.5 |
| OPA | ResNet50 | avgpool | 5/5 | late (final) | 0.6 | MiniLM | sentence | N/A | N/A | 1.0 |
| PPA | ResNet50 | avgpool | 5/5 | late (final) | 0.9 | MiniLM | sentence | N/A | N/A | 1.0 |
| RSC | ResNet50 | avgpool | 5/5 | late (final) | 0.8 | MPNet | sentence | N/A | N/A | 0.5 |

**Supplementary Table 2. Best vision DNN and LLM layers selected for each ROI: NSD (scenes).** Stability is the proportion of the 10 cross-validation splits in which the reported layer was selected.

| ROI | Vision DNN | Layer | Position | Depth | Stability | LLM | Layer | Position | Depth | Stability |
|---|---|---|---|---|---|---|---|---|---|---|
| V1 | VGG16 | pool3 | 3/8 | mid | 0.7 | MPNet | sentence | N/A | N/A | 0.6 |
| V2 | VGG16 | pool3 | 3/8 | mid | 1.0 | MPNet | sentence | N/A | N/A | 0.8 |
| V3 | VGG16 | pool4 | 4/8 | mid | 0.7 | BERT | L07 | 8/12 | mid | 0.5 |
| hV4 | VGG16 | pool4 | 4/8 | mid | 1.0 | BERT | L07 | 8/12 | mid | 0.4 |
| EBA | AlexNet | fc6 | 6/8 | late | 0.4 | GPT2 | h6 | 7/12 | mid | 0.9 |
| FFA | AlexNet | fc7 | 7/8 | late | 0.6 | GPT2 | h6 | 7/12 | mid | 0.9 |
| OFA | VGG16 | pool5 | 5/8 | mid | 1.0 | BERT | L07 | 8/12 | mid | 0.5 |
| OPA | VGG16 | pool5 | 5/8 | mid | 0.7 | GPT2 | h6 | 7/12 | mid | 0.8 |
| PPA | AlexNet | fc6 | 6/8 | late | 0.8 | BERT | L06 | 7/12 | mid | 0.4 |
| RSC | AlexNet | fc7 | 7/8 | late | 0.5 | MPNet | sentence | N/A | N/A | 0.4 |

**Supplementary Table 3. Best vision DNN and LLM layers selected for each ROI: BMD (dynamic events).** Stability is the proportion of the 10 cross-validation splits in which the reported layer was selected.

| ROI | Vision DNN | Layer | Position | Depth | Stability | LLM | Layer | Position | Depth | Stability |
|---|---|---|---|---|---|---|---|---|---|---|
| V1 | TimeSformer-divST | blk03 | 4/12 | early | 1.0 | MiniLM | sentence | N/A | N/A | 0.8 |
| V2 | TimeSformer-divST | blk03 | 4/12 | early | 0.9 | MiniLM | sentence | N/A | N/A | 0.6 |
| V3 | TimeSformer-divST | blk03 | 4/12 | early | 0.8 | MPNet | sentence | N/A | N/A | 0.5 |
| hV4 | tsm-r50 | stage4 | 4/4 | late (final) | 1.0 | MiniLM | sentence | N/A | N/A | 1.0 |
| EBA | tsm-r50 | stage4 | 4/4 | late (final) | 0.9 | MiniLM | sentence | N/A | N/A | 0.5 |
| FFA | tsm-r50 | stage4 | 4/4 | late (final) | 1.0 | MiniLM | sentence | N/A | N/A | 1.0 |
| OFA | tsm-r50 | stage4 | 4/4 | late (final) | 1.0 | MiniLM | sentence | N/A | N/A | 0.8 |
| OPA | VideoMAE-B | blk08 | 9/12 | late | 0.5 | MiniLM | sentence | N/A | N/A | 0.8 |
| PPA | VideoMAE-B | blk08 | 9/12 | late | 0.5 | MiniLM | sentence | N/A | N/A | 0.8 |
| RSC | VideoMAE-B | blk07 | 8/12 | mid | 0.5 | MiniLM | sentence | N/A | N/A | 0.8 |

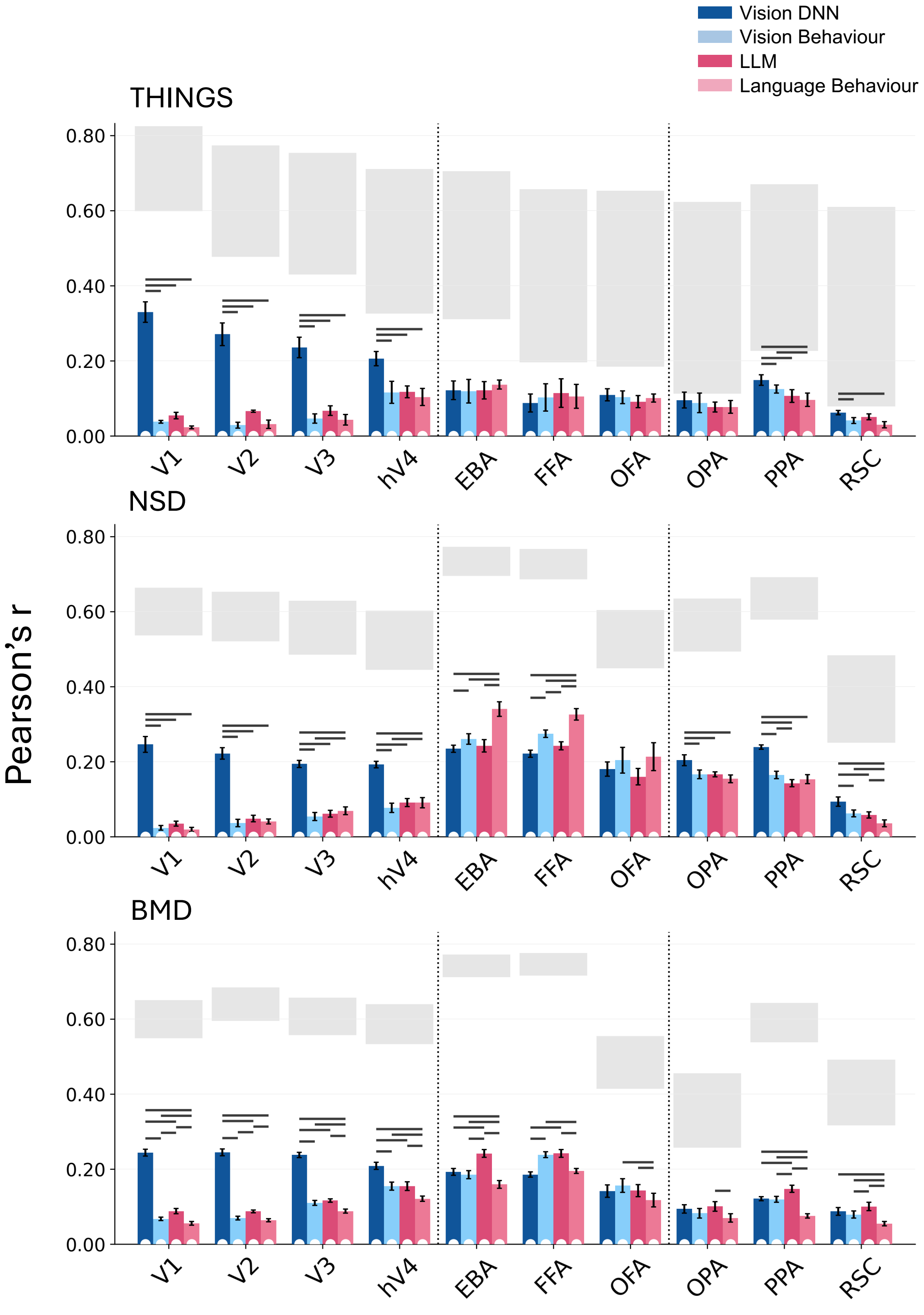


**Supplementary Figure 6. Brain alignment of vision and language predictors relative to the noise ceiling.** Coloured bars show the alignment values reported in Figures 4 and 5 (top panels) of the main text, presented together for all four predictors (mean ± SEM across participants). White dew-drops at the base of each bar indicate alignment significantly above zero, and dark grey horizontal lines indicate significant differences between predictors. Significance was assessed using two-sided Wilcoxon signed-rank tests for NSD and BMD and subject-level two-sided stimulus-label permutation tests for THINGS. All $p$-values were FDR-corrected across ten ROIs and were indicated as significant when $p < 0.05$. Light grey bars indicate the noise ceiling (Nili et al., 2014), which estimates the range of achievable alignment given inter-participant variability. The lower bound was the mean leave-one-out correlation between each participant's RDM and the average RDM of the other participants. The upper bound is the mean correlation between each participant's RDM and the group-mean RDM (including that participant).

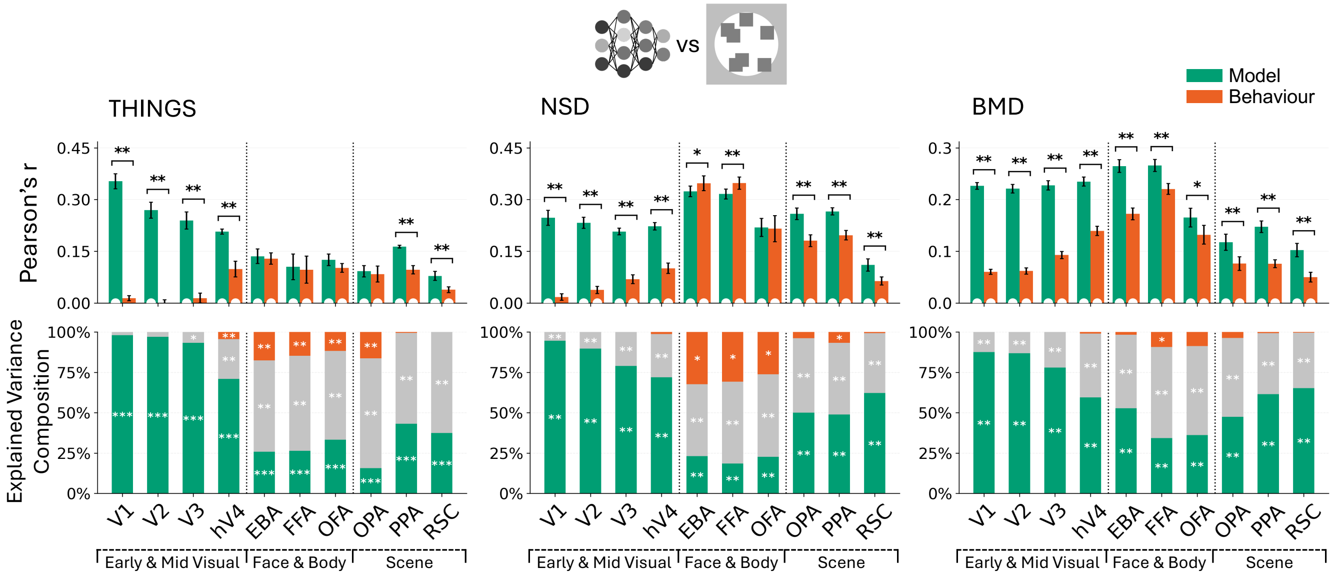


**Supplementary Figure 7. Behaviour-model comparison of brain alignment and variance partitioning.** Top: Alignment between joint behaviour and joint model RDMs and brain RDMs across ROIs for each dataset. The behaviour predictor is a non-negative weighted combination of the vision- and language-based similarity-judgement RDMs, and the model predictor is a non-negative weighted combination of the vision DNN and LLM RDMs across ROIs (layers selected per ROI by nested cross-validation). White dew-drop at the base of each bar indicates alignment significantly greater than zero. Brackets with asterisks denote significant differences between behaviour and model. Significance was assessed with two-sided Wilcoxon signed-rank tests for NSD and BMD, and subject-level two-sided stimulus-label permutation tests for THINGS. Bottom: Variance explained uniquely by behaviour, uniquely by model, and shared between behaviour and model, as a percentage of total explained variance across ROIs for each dataset. All $p$-values are FDR-corrected across ten ROIs. Significance: *$p < 0.05$, **$p < 0.01$, ***$p < 0.001$.

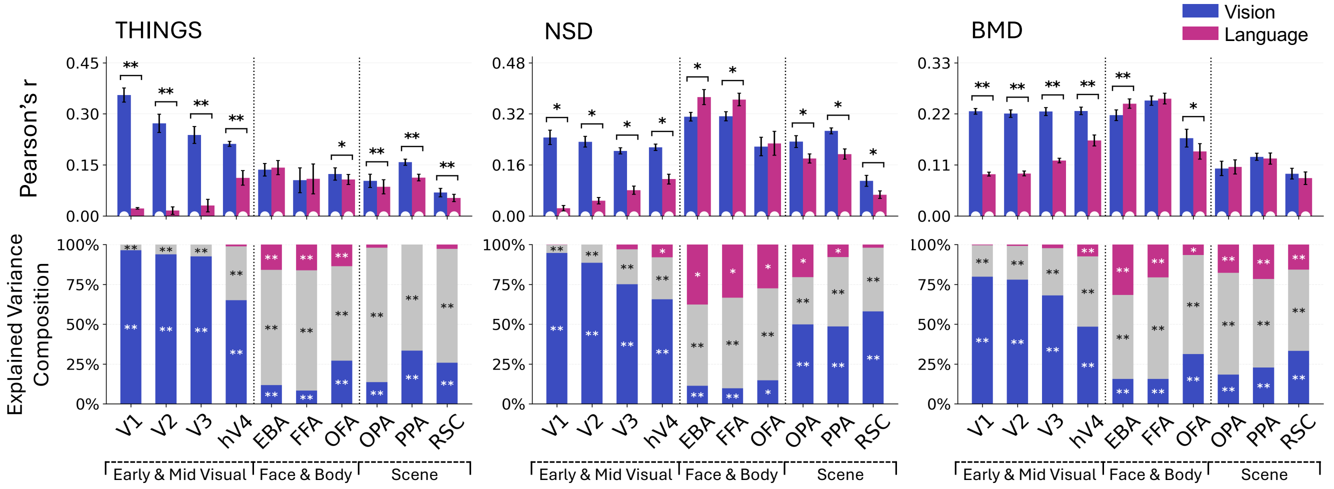


**Supplementary Figure 8. Brain alignment and variance partitioning of vision and language predictors.** Top: Alignment of vision- and language-derived RDMs with brain RDMs across ROIs and datasets. The vision RDM combines vision DNNs and vision-based similarity judgements, whereas the language RDM combines LLMs and language-based similarity judgements. White dew-drops at the base of each bar indicate alignment significantly above zero, and brackets with asterisks indicate significant differences between vision and language. Bottom: Variance explained uniquely by vision, uniquely by language, and shared between them, expressed as percentages of the sum of non-negative variance components for each ROI and dataset. Significance was assessed using two-sided Wilcoxon signed-rank tests for NSD and BMD and subject-level two-sided stimulus-label permutation tests for THINGS. Asterisks are shown only in segments of at least 3% of their bar for display purposes. Three significant positive components fall below this threshold, all representing unique variance explained by language: THINGS hV4 (1.1%), NSD RSC (2.0%) and BMD V3 (2.3%). All $p$-values were FDR-corrected across ten ROIs. Significance levels: *$p < 0.05$, **$p < 0.01$, ***$p < 0.001$.
